# Cryo-electron tomography reveals the multiplex anatomy of condensed native chromatin and its unfolding by histone citrullination

**DOI:** 10.1101/2022.07.11.499179

**Authors:** Nathan Jentink, Carson Purnell, Brianna Kable, Matthew Swulius, Sergei A. Grigoryev

**Affiliations:** Penn State University College of Medicine, Dept. Biochemistry & Molecular Biology, H171, Milton S. Hershey Medical Center, P.O. Box 850, 500 University Drive, Hershey, PA 17033

## Abstract

Nucleosome chains fold and self-associate to form higher order structures whose internal organization is unknown. Here, cryo-electron tomography (cryo-ET) of native human chromatin reveals novel folding motifs such as 1) non-uniform nucleosome stacking, 2) intermittent parallel and perpendicular orientations of adjacent nucleosome planes, and 3) an inverse zigzag nucleosome chain path, which deviates from the direct zigzag topology seen in reconstituted nucleosomal arrays. By examining these self-associated structures, we observed prominent nucleosome stacking *in-cis* and anti-parallel nucleosome interactions *in-trans,* which are consistent with partial nucleosome interdigitation. Histone citrullination strongly inhibits nucleosome stacking and self-association with a modest effect on chromatin folding, while the reconstituted arrays showed a zigzag topology which undergoes a dramatic unfolding induced by histone citrullination. This study sheds light on the internal structure of compact chromatin nanoparticles and suggests a novel mechanism for how epigenetic changes in chromatin are retained across both open and condensed forms of chromatin.

## INTRODUCTION

The DNA in eukaryotic chromatin is repeatedly coiled around histone proteins, forming arrays of 10 nm nucleosomes. Each nucleosome contains a core of about 147 bp of DNA that makes approximately 1.7 left superhelical turns around an octamer of four pairs of core-histones H2A, H2B, H3, and H4 (Richmond and Davey, 2003). Nucleosome “beads” are connected by extended linker DNA “strings” (10 to 100 bp in length) into “beads-on-a-string” nucleosome chains. These nucleosome chains are packed through chromatin higher-order folding mechanisms to achieve a 400–1,000-fold compaction of DNA in the relatively decondensed interphase chromatin (Hu et al., 2009) and 10,000-fold compaction in condensed metaphase chromosomes (Alberts et al., 2015). Chromatin higher-order folding limits DNA accessibility for transcription factors (Iwafuchi-Doi and Zaret, 2014; Poirier et al., 2008) and DNA repair machinery (Supek and Lehner, 2015) during interphase, and mediates chromosomal integrity during cell division (Schneider et al., 2021). Solving the 3D organization of native nucleosome arrays in their condensed states, and understanding the molecular mechanism driving tight nucleosome packing, would bring about a fundamental advance in understanding the processes underlying epigenetic gene regulation and chromosomal stability.

Chromatin higher-order folding has been suggested to comprise a hierarchy of structural levels. The nucleosome arrays (the primary structural level) first fold longitudinally into the 30-nm chromatin fibers (the secondary structural level) and then the fibers would associate latitudinally to form self-associated tertiary structures that may further expand to form condensed chromatin or chromosomes (Woodcock and Dimitrov, 2001). 30-nm chromatin fibers have been observed by *in-situ* EM in the nuclei of some terminally-differentiated cells, in nucleated chicken erythrocytes (Scheffer et al., 2011), starfish sperm (Horowitz et al., 1994), and mouse retina (Kizilyaprak et al., 2010). They have also been reconstituted in multiple experiments that employed arrays of artificial nucleosome-positioning sequences which showed a prominent secondary level folding along the two-start zig-zag (Bednar et al., 1998; Dorigo et al., 2004; Garcia-Saez et al., 2018; Routh et al., 2008; Schalch et al., 2005; Song et al., 2014). Despite many years of intensive studies, however, the existence of 30-nm fibers, or any regularly folded structures above the primary nucleosome level, remains elusive in most eukaryotic cells (Grigoryev, 2018; Joti et al., 2012; Luger et al., 2012). Neither electron microscopy (Cai et al., 2018b; Eltsov et al., 2008; Fussner et al., 2011; Ou et al., 2017) nor super-resolution fluorescence imaging (Nozaki et al., 2017; Ricci et al., 2015) have shown any extended chromatin secondary structures above the 10-nm diameter. Moreover, the tertiary-level nucleosome condensates induced by Mg^2+^ did not show any distinct secondary structures as their intermediates (Maeshima et al., 2016). These nucleosome condensates appeared to be remarkably fluid and were proposed to be formed by the liquid-liquid phase separation (LLPS) (Gibson et al., 2019) forming either liquid droplets or semi-solid hydrogels, depending on the experimental conditions and the state of histone modification (Strickfaden et al., 2020).

Nevertheless, capturing nucleosome interactions in mammalian cell nuclei (Grigoryev et al., 2016; Hsieh et al., 2020; Krietenstein et al., 2020; Risca et al., 2017) showed a distinct pattern of nucleosome proximities consistent with the two-start zig-zag folding. Furthermore, short nucleosome clusters, or nanodomains, were observed by super-resolution microscopy (Fang et al., 2018; Kieffer-Kwon et al., 2017; Ricci et al., 2015), and some higher-order folds, loops, and hubs formed by closely juxtaposed nucleosomes were observed by electron tomography (Ou et al., 2017) indicating that, despite the absence of the long regular fibers, some discrete elements of secondary and tertiary structure may underlie the multiplex chromatin folding in living cells.

Cryo-electron tomography (Cryo-ET) allows one to resolve biological structures embedded in thin layers of vitrified ice or in thinned cellular sections at nanoscale resolution (Hylton and Swulius, 2021; Saibil, 2022). Unlike a single-particle cryo-EM approach, which can achieve angstrom scale resolution by averaging images of many thousand particles, Cryo-ET is used to resolve individual molecules and molecular assemblies, which makes it ideal for analysis of multiplex nucleosome chain conformations. Cryo-ET was previously used to resolve nucleosome cores within cellular sections (Cai et al., 2018a; Eltsov et al., 2018; Mahamid et al., 2016; Xu et al., 2021) as well as *in situ*-crosslinked and isolated chromatin (Arimura et al., 2021), though the nucleosome chain path and nucleosome interaction patterns have not been resolved. With isolated native interphase chromatin and metaphase chromosomes, the linker DNA can be resolved for long nucleosome arrays unfolded at relatively low ionic strength (Bednar et al., 1998; Beel et al., 2021; Grigoryev et al., 1999).

Here we applied Cryo-ET to analyze condensed native nucleosome arrays isolated from human cells and embedded in vitrified ice without any fixation or staining. Using Cryo-ET in combination with nanoscale stereological modeling of individual nucleosomes, we traced consecutive nucleosomes within the condensed native human chromatin and determined its key spatial parameters such as nucleosome center-to-center distances and plane angles. Within the most condensed tertiary structures, we observed abundant parallel nucleosome stacking *in-cis* and also anti-parallel nucleosome interaction *in-trans,* consistent with partial nucleosome interdigitation. We have shown that histone citrullination by PAD4, which causes massive chromatin unfolding during NETosis (Brinkmann and Zychlinsky, 2012; Wang et al., 2009) and contributes to deep vein thrombosis (Martinod et al., 2013), cancer metastasis (Yang et al., 2020), and COVID-19 pathology (Barnes et al., 2020; Veras et al., 2020), causes a dramatic unfolding of the secondary and tertiary higher-order structures by disrupting the nucleosome stacking interactions. Furthermore, by comparison with reconstituted nucleosome arrays, we were able to deduce secondary structural features and linker DNA conformation specific for the native chromatin that explain its folding into discrete nanoparticles, in contrast to the extended zigzag 30 nm fiber in the reconstituted nucleosome arrays. We conclude that cryo-ET in combination with stereological nucleosome modeling is a powerful experimental approach that can reveal nanoscale chromatin structural transitions vital for understanding the mechanism(s) of gene regulation and NETosis and provides a valuable resource for imaging and analysis of the multiplex nucleosome chain conformations in living cells.

## RESULTS

### Cryo-ET reveals secondary and tertiary structures in Mg^2+^-condensed human chromatin

We isolated MNase-fragmented soluble chromatin from human K562 cells, and fractionated chromatin by sucrose gradient ultracentrifugation to select a chromatin fraction containing ∼ 12 nucleosomes per particle by DNA size (Fig. 1 A), which is the same size as the 12-mer repeats used in most experiments with reconstituted chromatin. We induced chromatin condensation by the divalent cation, Mg^2+^, which causes full compaction of nucleosome arrays into zig zag fibers concentrations 0.5 - 1 mM (Dorigo et al., 2004; Garcia-Saez et al., 2018) which are within the physiological range of free Mg^2+^ *in vivo* (Maeshima et al., 2018). Upon increased Mg^2+^, chromatin fractions showed a sharp self-association around 1.4 mM without a significant difference between two biological replicates (Fig. 1 B) and very similar to the Mg^2+^-dependent self-association of nucleosome arrays into the bulky tertiary structure described previously (Carruthers et al., 1998; Maeshima et al., 2016). To monitor the process of chromatin condensation in solution, we crosslinked chromatin fractions with glutaraldehyde at different Mg^2+^ concentrations and resolved the chromatin particles on a native DNP (DNA/protein) agarose gel (Fig. 1 C). At 0.75 mM, the chromatin particles remained homogeneous with a notable increase in mobility indicating formation of a condensed secondary structure. At 1.0 mM, a retarded band appeared indicating partial self-association. This self-association gradually increased with increasing Mg^2+^ concentration until no material entered the agarose gel at 2 mM Mg^2+^, which we considered to be the final point for the tertiary structure formation in our experiments. Similarly fixed samples were examined by negative stain transmission EM and were consistent with the DNP gel analysis showing individual condensed nucleosome arrays at 0.75 mM Mg^2+^ and bulky nucleosome condensates at 1.25 and 2.0 mM Mg^2+^ (Fig. 1 D).

**Figure 1:**
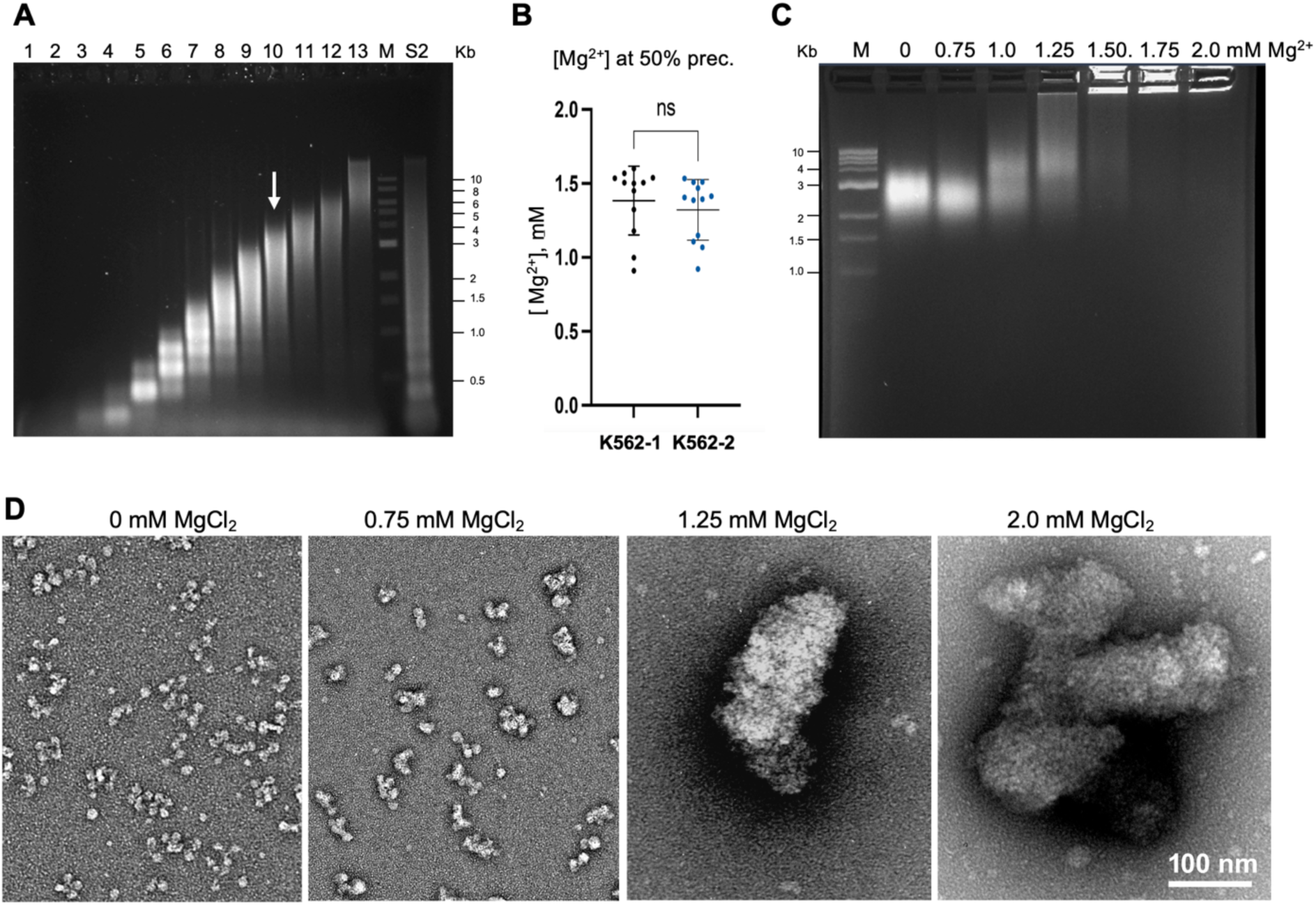
Biochemical and TEM characterization of Mg^2+^-dependent secondary and tertiary structures in native human chromatin. (A) Agarose gel shows DNA of native chromatin isolated from K562 cells and fractionated by ultracentrifugation of sucrose gradients (lanes 1-13), DNA size markers (lane 14 (M)), and total soluble chromatin (lane S2). White arrow indicates the chromatin fraction #10 selected for TEM and Cryo-ET imaging. (B) Mg^2+^-induced 50% precipitation points determined for chromatin fractions #10 isolated from two independent batches of K562 cells. (C) DNP agarose gel showing native electrophoresis of the K562 chromatin crosslinked by glutaraldehyde at the indicated concentration of Mg^2+^. Lane M: DNA size markers. (D) TEM of the K562 chromatin crosslinked by glutaraldehyde at the indicated concentration of Mg^2+^ shows compaction of individual particles at 0.75 mM Mg^2+^ and formation of bulky self-associates at 1.25 and 2.0 mM Mg^2+^. Scale bar: 100 nm.

We then examined native chromatin particles vitrified in HNE buffer (5 mM NaCl, 0.1 mM EDTA, 10 mM HEPES-NaOH, pH=7.5) with and without increasing Mg^2+^ concentrations. In the low-salt HNE buffer without Mg^2+^ we observed unfolded nucleosome chains (Fig. 2 A) very similar to our previous Cryo-EM of native chromatin (Bednar et al., 1998; Grigoryev et al., 1999). In the presence of 0.75 mM Mg^2+^, however, the chromatin showed compaction of nucleosome disks into flat ladder-like nucleosome assemblies (Fig. 2 B) resembling those previously observed by single-particle Cryo-EM of Mg^2+^-condensed reconstituted clone 601-nucleosome arrays (Garcia-Saez et al., 2018). We did not observe helical fibers like those observed by glutaraldehyde fixation (Song et al., 2014). Unlike reconstituted clone 601 chromatin, the native nucleosome arrays were highly heterogeneous, with most containing stacks of several nucleosomes interrupted with unstacked nucleosome cores, and some arrays contained no nucleosome stacking at all. Chromatin vitrified at 1.25 mM and 2 mM Mg^2+^ showed prominent self-association in which separate nucleosome arrays merged together to form bulky tertiary structures (Fig. 2 C, D). Under a lower magnification, we observed large electron-dense chromatin clusters with diameters often exceeding 1 μm (Fig. 2 E). The tertiary structures formed in the presence of either 1.25 or 2 mM Mg^2+^ were rapidly flattened in the process of vitrification and were amenable to Cryo-ET imaging in a completely vitrified state. Though we could not resolve separate nucleosome arrays within the tertiary structures, the nucleosome packing retained a high degree of heterogeneity with apparent stacks of several nucleosomes (seen as edge-to edge) and regions where no stacking was observed. While we routinely used 20 min. periods of Mg^2+^ incubation, in parallel control experiments, we examined chromatin particles incubated for 48 hours in the presence of Mg^2+^ and BSA-covered EM fiduciary gold particles by SDS-PAGE to show a complete absence of histone degradation (Suppl. Fig. S1 A). Additionally, we did not observe any changes in chromatin condensation upon incubation for 3 and 16 hrs. by cryo-ET (Suppl. Fig. S1 B, C). The chromatin samples, vitrification conditions, and resulting cryotomograms are listed in the Suppl. Table 1.

**Figure 2:**
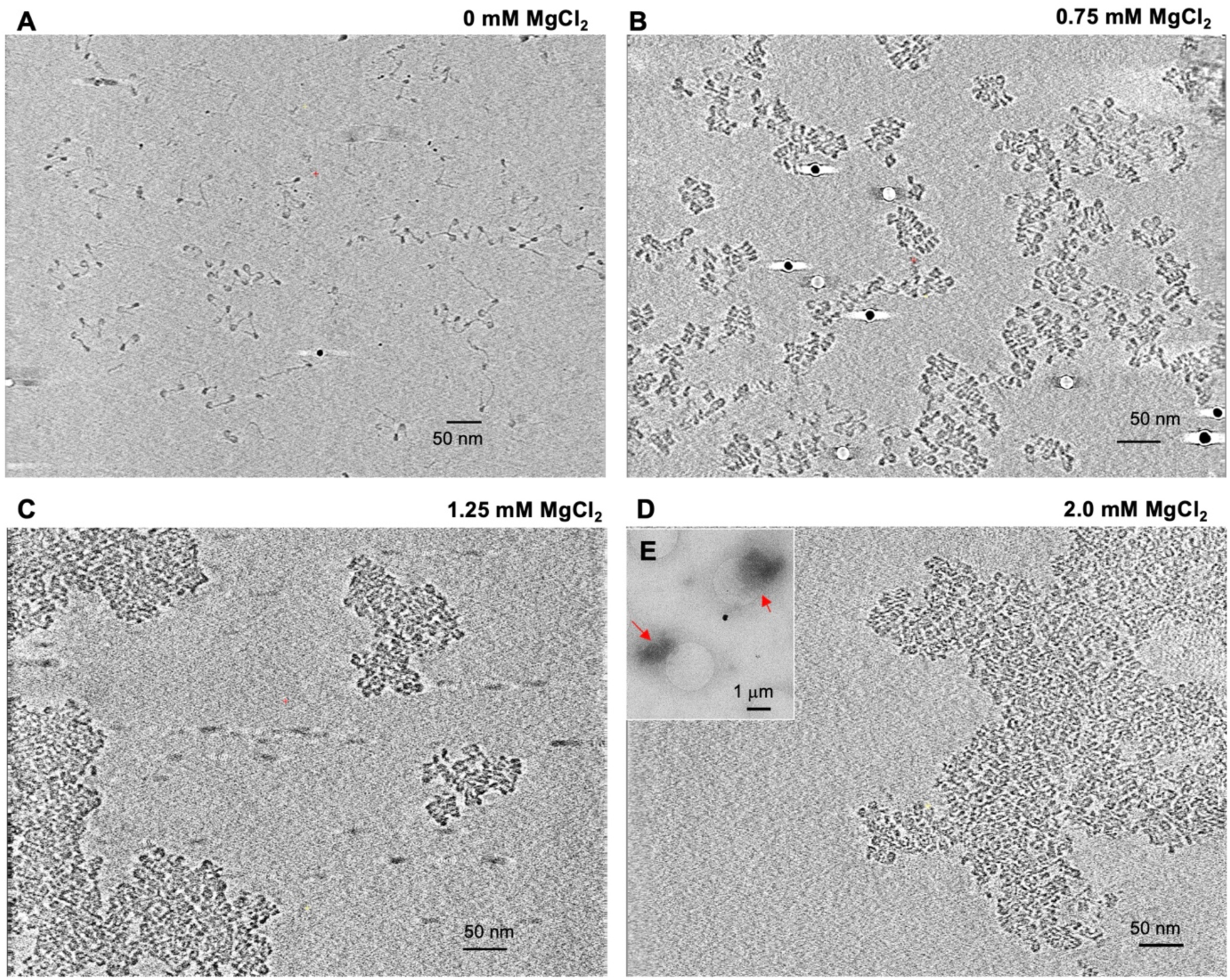
Cryo-ET reveals abundant nucleosome stacking within the secondary and tertiary chromatin structures. (A-D) Representative Z-series slices from Cryo-ET tomograms of K562 chromatin embedded in vitrified ice at the indicated concentration of Mg^2+^ together with 10 nm fiduciary gold particles show open nucleosome arrays in HNE buffer without Mg^2+^ (A, TS_1_3), compaction of individual particles at 0.75 mM Mg^2+^ (B, TS_4_4), and formation of bulky self-associates at 1.25 (C, TS_6_1) and 2.0 mM Mg^2+^(D, TS_8_2). Mag: 53,000-x; scale bar 50 nm. (E) an insert showing a portion of direct Cryo-EM image (search mode) of K562 chromatin embedded in vitrified ice at 2.0 Mg^2+^. Mag: 2,250-x; scale bar: 1 μm.

Concerned that our observations may be due to our use of proliferating K562 cells, many of which could be affected by the chromatin-condensing factors activated in the metaphase of cell division, we analyzed chromatin from cells blocked either in the G1-phase by mimosine or in the S-phase by hydroxyurea (Suppl. Fig. S2 A-C). For all cases, we observed similar Mg^2+^ -dependent self-association (Suppl. Fig. S2 D, E) and the folding of nucleosome arrays at 0, 0.75, and 2.0 mM Mg^2+^ was indistinguishable from that of control chromatin by Cryo-EM (cf. Fig 2 and Suppl. Fig. S2 F, G), showing that the extent of nucleosome stacking does not vary with the cell cycle. Furthermore, we analyzed chromatin from two other vertebrate cell types – human HeLa cells with a predominantly open euchromatin (Grigoryev et al., 2016) and mouse retina cells at postnatal stage 21 which have completely exited the cell cycle and have acquired abundant condensed heterochromatin (Popova et al., 2013). Since previous experiments with reconstituted chromatin showed that nucleosome repeat length (NRL) was an important factor regulating both secondary (Correll et al., 2012) and tertiary structure in the presence of Mg^2+^ (Gibson et al., 2019) we analyzed the NRL associated with these three cell types. (Suppl. Fig. S3 A-C) finding the NRL in Hela (184 bp) to be shorter than in K562 (190 bp) and mouse retina (194 bp). While HeLa chromatin showed a slightly higher mid-transition point of self-association (Suppl. Fig. S3 D), by Cryo-ET, all three cell types underwent very similar compaction resulting in abundant nucleosome juxtaposition at 0.75 mM Mg and large nucleosome condensates with heterogeneous nucleosome stacking at 2 mM Mg (cf. Fig 2 and Suppl. Fig. S3 E,F). We concluded that the major effect of physiological Mg^2+^ on chromatin structure is promoting stacking of nucleosome disks and that the tertiary structures of condensed native chromatin are universal and independent of the cell cycle. Since K562 chromatin showed the most prominent nucleosome stacking in our preliminary experiments, it was chosen for further detailed analysis by Cryo-ET and nanoscale stereological modeling.

### Spatial modeling and stereological nanoscale analysis of the secondary chromatin structure

In our cryotomograms, under all conditions, individual nucleosome cores were clearly resolved. DNA linkers were fully resolved in the uncondensed chromatin, and in thin ice we could observe additional fine structures connecting the nucleosome cores and linkers in the unfolded chromatin (Fig 3A). These are most likely the long N-terminal tail of histone H3 and C-terminal tail of linker histone (white arrows on Fig. 3 A) as well as bulkier protrusions emanating from the nucleosome cores (arrowhead on Fig. 3 A).

**Figure 3:**
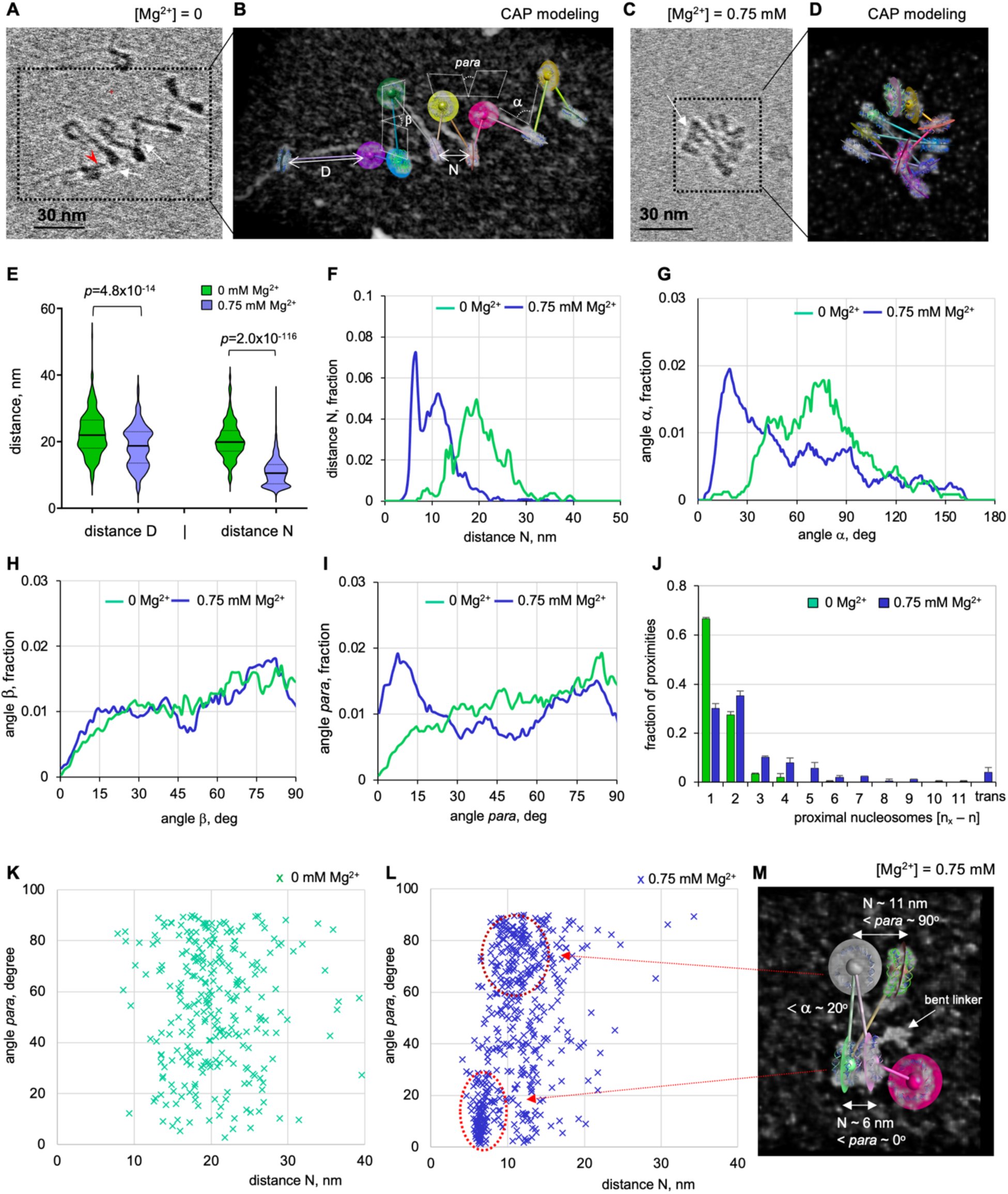
Cryo-ET and stereological modeling reveal the key parameters of nucleosome chain folding in Mg^2+^-condensed native chromatin. (A) A cropped Cryo-ET tomogram (TS_1_1) of the K562 chromatin vitrified in the low-salt HNE buffer without Mg^2+^ ion. In addition to the apparent nucleosomes, the positions of putative histone tails are indicated by white arrows and a bulky extranucleosomal protrusion by the red arrowhead. (B) The cropped image shown in (A) was processed to generate a CAP model (CAP #1_1_1). The scheme shows variables measured from the CAP model: distances D and N and angles α, β, and *para*. (C) A cropped Cryo-ET tomogram of the K562 chromatin vitrified at 0.75 mM Mg^2+^(TS_4_1). Stacks of parallelly oriented nucleosome disks are indicated by white arrows. (D) The image of Mg^2+^-condensed chromatin shown in C was processed to generate a CAP model (CAP #4_1_1). (E–I) Violin plot (E) and frequency distribution profiles of distances D (panel E), N (E, F), and angles α (G), β (H) and *para* (I) obtained for arrays of nucleosomes vitrified at 0 mM Mg^2+^ (green curves, n = 256 (D), 270 (N), 220 (α), 256 (β), 270 (*para*)) and arrays of nucleosomes vitrified at 0.75 mM Mg^2+^ (blue curves, n = 402 (D), 507 (N), 326 (α), 396 (β), 507 (*para*)). (J) Distribution of pairwise nucleosome proximities determined for arrays of nucleosomes vitrified at 0 mM Mg^2+^ (green columns, n = 267) and arrays of nucleosomes vitrified at 0.75 mM Mg^2+^ (blue columns, n = 345). (K, L) Two-dimensional plot showing distributions of the internucleosomal angle *para* vs. the distance N for nucleosomes vitrified at 0 mM Mg^2+^ (K) and 0.75 mM Mg^2+^ (L). (M) A cropped image of Mg^2+^-condensed chromatin (TS_3_1) was processed to generate a CAP model (CAP #3_1_9). Angles *para* and distances N between selected nucleosome pairs as well as one bent linker DNA are indicated.

For quantitative stereological analysis of chromatin folding, we built nucleosome chain models by cropping individual nucleosome array subvolumes from both 0 and 0.75 mM Mg^2+^ cryotomograms, and rigid-body fitting PDB models of the nucleosome core (PDB2CV5) into the density maps. The centroid and plane of each nucleosome was calculated and connected into a chain by axes (Fig. 3 B and D) to form a centroid/axes/plane (CAP) model for each nucleosomal array. In condensed chromatin, the nucleosome cores were resolved, but it was difficult to trace all linker DNA and their conformations, so ∼21% of linkers in the condensed chromatin were unaccounted for in our modeling. Snapshots of individual CAP models are shown on Suppl. Fig. 4 A and 4 B.

To perform stereological analysis, we modified the two-angle model describing chromatin zig zag folding (Grigoryev, 2018) and recorded five values for each nucleosome: 1) the center-to-center distance D from the next nucleosome in the chain, 2) the center-to-center distance N from the nearest nucleosome in 3D space, 3) the angle α between the two axes connecting each set of three consecutive nucleosomes in a chain, 4) the angle β between the planes of consecutive nucleosome pairs, and 5) the angle *para* between the planes of each nucleosome and its nearest neighbor in the 3D space (Fig 3 B). It is important to note that the nearest neighbor nucleosome distance N and angle *para* were completely monitored for all open and condensed nucleosomes, providing the most robust data to monitor chromatin secondary structural compaction. Comparison between the measurements for two independent batches of K562 cells showed a strong similarity between the nucleosome folding parameters (Suppl. Fig. S5). All our distance and angle measurements for individual nucleosome arrays are included in the Suppl. Table 2.

Modeling and comparative analysis of multiple nucleosomes vitrified at 0 mM Mg^2+^ (n = 268) and 0.75 mM Mg^2+^ (n = 486) showed that the distance between consecutive nucleosomes (D) was reduced rather modestly, from 22.4 to 18.6 nm on average, *p*<10^-13^ (see Fig. 3 E and Suppl. Fig. S6 A). This is consistent with partial linker DNA bending by Mg^2+^, as predicted by the heteromorphic zigzag model (Grigoryev et al., 2016), but clearly inconsistent with the overall bending of linker DNA in the solenoidal models (Finch and Klug, 1976). An example of partial linker DNA bending is shown in Fig. 3 M. In comparison, the distance N between the nearest neighbor nucleosomes was reduced much more profoundly, from 20.5 to 10.9 nm average, *p*<10^-115^, producing a sharp peak at 6 nm (Fig. 3 F), which corresponds to the average distance between parallel stacked nucleosomes. There was also a broader peak at 9-12 nm which corresponds to the distance between juxtaposed, but perpendicularly oriented, nucleosome disks (Fig. 3 F). We defined two nucleosomes as stacked if their distance N was less than 8 nm and calculated that 30.5% of all nucleosomes vitrified at 0.75 mM Mg^2+^ were stacked (no stacking occurred without Mg^2+^). The distribution of stacked nucleosomes among individual nucleosome arrays vitrified at 0.75 mM Mg^2+^ varies widely, from 0 to 78 % of stacked nucleosomes per particle (Suppl. Fig. S6 H), reflecting the high structural heterogeneity of the native chromatin observed by Cryo-ET (Fig. 2).

Plotting the distribution of angle α shows a sharp increase in values below 40°, in condensed chromatin, generating a prominent peak at ∼20° (Fig. 3 G and Suppl. Fig. S6 B). That being said, the total average angle α values did not change between conditions as significantly as the nucleosome distances (*p*<10^-6^) reflecting the wide distribution of angle α in the folded and the unfolded states (Fig. 3 G). When we compared only the lowest 20% of angles α, however, we observed a much stronger difference with *p*<10^-25^, and the top 20 % were statistically identical (Suppl. Fig. S6 B). This observation is consistent with the heterogeneous angle α populations, and likely reflect a variable association with linker histone H1, the chromatin protein that reduces the angles between linker DNA upon binding near the nucleosome dyad axis (Bednar et al., 2017; Zhou et al., 2015).

In contrast to the angle α distribution, angle β does not change significantly in the Mg^2+^-condensed chromatin (Fig. 3 H and Suppl. Fig. S6 C), suggesting that the heteromorphic nucleosome orientations resulting from variable linker DNA length (Woodcock et al., 1993) are not altered during compaction, generating the strikingly nonuniform stacking observed in our native chromatin by Cryo-ET (Fig. 3, panels D and M.). This is in sharp contrast to what was observed for the Mg^2+^-condensed reconstituted nucleosome arrays with regular nucleosome linkers (Garcia-Saez et al., 2018).

In Mg^2+^-condensed chromatin, the frequency of angle *para* displayed a prominent increase at values below 20° with a broad peak at ∼10° (Fig. 3 I and Suppl. Fig. S6 D), consistent with the parallel nucleosome stacking observed in our cryotomograms. Like with angle α, the change in average angle *para* was only modestly significant (*p*<10^-5^) but became highly significant (p<10^-25^) when only the lowest 20% of angles *para* were compared (Suppl. Fig. S6 D).

To look for associations between the nucleosome plane angle and the internucleosomal distance, we plotted distributions of nucleosome angles para versus the corresponding distance between neighboring nucleosomes (N). The two-dimensional plots reveal two distinct areas at 0.75 mM Mg^2+^(Fig. 3 L) that are absent at 0 mM (Fig. 3 K). One is below 25° and is centered at the distance N ∼ 6nm, which corresponds to the tightly stacked parallel nucleosome disks. The other, more dispersed range at 60-90° angle, is centered at the distance N ∼ 9-12 nm and corresponds to perpendicular nucleosome disks (Fig. 3 M). Overall, the parallel nucleosome disk stacking, though abundant in native K562 chromatin, is non-uniform and is interspersed with nucleosomes juxtaposed almost perpendicularly towards each other.

Finally, to compare the chromatin structures resolved by Cryo-ET with our previous nucleosome interaction map *in situ* (Grigoryev et al., 2016), we measured the nucleosome pairwise proximities (i±n). This was done by numbering the consecutive nucleosomes in the arrays and calculating the absolute difference (n) between the number of each nucleosome in the chain (i) and the number of its closest counterpart (k), see Suppl. Table 2. In the unfolded state, the majority of nucleosomes had their most proximal neighbors at nucleosome i±1 expected for an extended nucleosome chain with a minority of nucleosome proximities at i±2 corresponding to a partial zigzag folding similar to the unfolded chromatin isolated from uncrosslinked human cells tested by EMANIC. In contrast, the Mg^2+^-condensed native chromatin showed a strong decrease in the nucleosome proximities at i±1 together with a profound increase in i±2, i±3, and i±4 (Fig. 3 J). This pattern is strikingly similar to the one previously observed by EMANIC (Grigoryev et al., 2016), (Suppl. Fig. S6 G), and also consistent with the genome-wide nucleosome interaction mapping in mammalian cells (Hsieh et al., 2020; Krietenstein et al., 2020; Risca et al., 2017). We thus concluded that the overall secondary structure of native chromatin observed by Cryo-ET is highly heterogeneous and remarkably similar to that observed *in situ*.

### Chromatin condensates retain abundant nucleosome stacking in-cis and in-trans

For our analysis of the secondary chromatin structures (Fig. 3), we focused on clearly separate nucleosome arrays, but even at 0.75 mM Mg^2+^ some arrays formed inter-array contacts mediated by close nucleosome interactions *in-trans*. Examples of such interacting arrays and their CAP models are shown in Fig 4A (panels 1 – 6). Remarkably, in contrast to the nucleosome stacking *in cis* between nucleosomes with parallel-oriented dyad axes, the nucleosomes connecting different arrays were either stacked in an anti-parallel orientation or were only partially overlapping (red arrows on panels 1 – 6).

**Figure 4:**
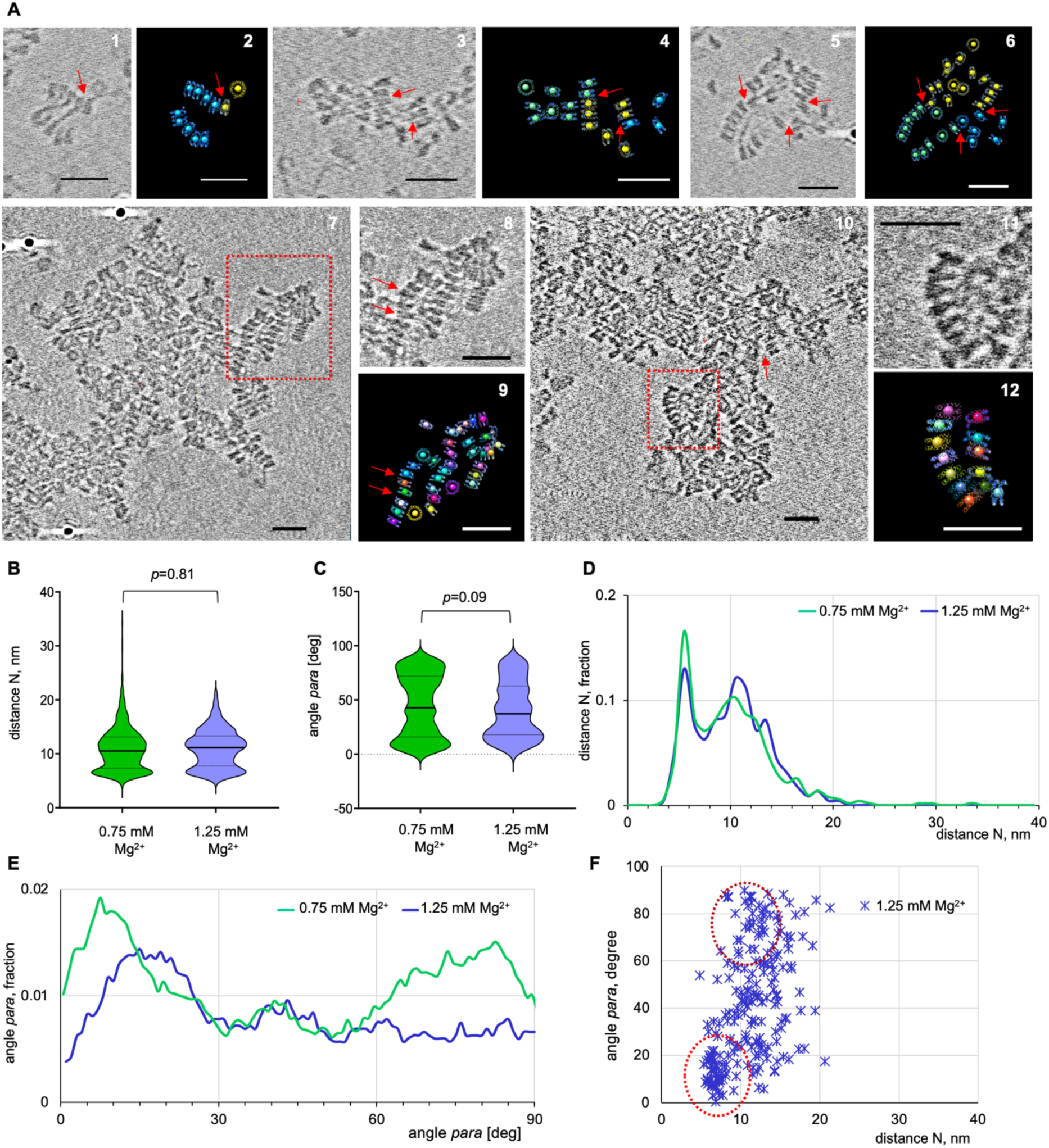
Cryo-ET and 3D modeling reveal *cis-* and *trans-*internucleosomal stacking within the self-associated native nucleosome arrays. (A) Panels 1, 3, and 5: cropped Cryo-ET tomograms showing K562 arrays vitrified at 0.75 mM Mg^2+^ and connected through nucleosome interactions *in-trans* (TS_ 3_3 (1 and 3) and #3_2 (5) . Panels 2, 4, and 6: the images in panels 1, 3, and 5 were processed to generate CAP models (CAP#3_3_1 (2), 3_3_3 (4), and 3_2_1 (5)). Panels 7, 8: Cryo-ET tomogram and a cropped area showing K562 nucleosome arrays vitrified at 1.0 mM Mg^2+^ (TS_5_2). Panels 9: the image in panels 8 was processed and modeled as described in the legend for Fig. 3 B (CAP#5_2_1). Panels 10, 11: Cryo-ET tomogram and a cropped area showing K562 nucleosome arrays vitrified at 1.25 mM Mg^2+^ (TS_6_2). Panels 12: the image in panels 11 was processed and modeled as described in the legend for Fig. 3 B (CAP#6_2_1). Scale bars: 30 nm. (B-E) Violin plots (B, C) and frequency distribution profiles (D, E) of distances N (B, D), and angle *para* (C, E) showing calculated for individual arrays of nucleosomes vitrified at 0.75 mM Mg^2+^ (green curves, n = 507) and cropped areas of nucleosome condensates vitrified at 1.25 mM Mg^2+^ (blue curves, n = 223). (F) Two-dimensional plot showing distribution of the internucleosomal angle *para* vs. the distance N for the nucleosomes vitrified at 1.25 mM Mg^2+^.

At 1.0 mM Mg^2+^, nucleosome arrays were more strongly associated, and they formed some bulky tertiary structures intermixed with detached nucleosome arrays (Fig 4A, panels 7–9, Suppl. Fig. S7, top row). These particles formed thin edges where the nucleosomes were self-associated two-dimensionally and therefore could be resolved by Cryo-ET for CAP modeling. A close-up of such an edge shows abundant nucleosome stacking *in-cis* with some nucleosomes stacked *in-trans*. Remarkably, some of these nucleosomes were engaged in a distinct structural motif in which the trans-interacting nucleosome disks were folded into stacks of antiparallel nucleosomes shown by red arrows in panels 8 and 9. Such a nucleosome arrangement is consistent with earlier nucleosome interdigitation models proposed for global chromatin condensation in heterochromatin (Grigoryev, 2004) and in interphase chromosomes (Chicano et al., 2019). However, in the Mg^2+^-condensed native chromatin the interdigitation involved small patches of 4 – 6 nucleosomes connecting distinct nucleosome arrays *in-trans* but not spreading globally to the majority of condensed chromatin.

At Mg^2+^ concentrations exceeding 1 mM, nucleosome arrays formed bulky condensates larger than 100-nm thick (Fig. 1 D), but during vitrification, these nucleosome condensates were flattened by surface tension to 50-80-nm thick, facilitating their imaging by cryo-ET (Fig 4A, panels 10–12, Suppl. Fig. S7, middle and bottom rows). Nucleosomes buried in the middle layers of the condensed chromatin particles showed distinct patterns of “mud-brick layering” formed by the nucleosome edges in the side projection at Y=90° (left panels in Suppl. Fig. S7). Our denoising methods did not yield clearly distinguishable individual nucleosomes in the middle of these large condensate particles, but we were able to resolve and model patches of condensed nucleosomes near the edge.

Stereological analysis of nucleosomes vitrified at 1.25 mM Mg^2+^ was compared to the nucleosomes vitrified at 0.75 mM Mg^2+^, and it showed that the average distance between the nearest neighbor nucleosomes (N) did not change significantly (*p* = 0.81, see Fig. 4 B). In both cases, the distribution of distances N showed a sharp peak at 6 nm, corresponding to the stacked nucleosomes, with a lower and broader peak at 9-12 nm (Fig. 4 D). In 1.25 mM Mg^2+^, the average value of angle *para* did not change significantly (Fig. 4 C), but the distribution did display a notable decrease at the values above 70° (Fig. 4 E). By comparing the two-dimensional plots of nucleosome angle para with the corresponding internucleosome distance (N), we observed a dissipation of the clustering at 60-90° α and the distance N ∼ 9-12 nm at 1.25 mM Mg^2+^ (the top dashed circle on Fig. 4 F), which corresponds to closely apposed but perpendicular nucleosome disks. The clustered area below 25° angle *para* and the distance N ∼ 6nm corresponding to the tightly stacked parallel nucleosome disks remained pronounced at 1.25 mM Mg^2+^ (bottom dashed circle; cf. Fig. 4 F and Fig. 3 L). Thus, in contrast to the previous lower resolution imaging suggesting the fully disordered nature of nucleosome condensates induced by Mg^2+^ (Gibson et al., 2019; Maeshima et al., 2016; Strickfaden et al., 2020), our cryo-EM images showed a clear retention of tight nucleosome stacking in the Mg^2+^-induced tertiary chromatin structure.

### PAD4-dependent histone citrullination inhibits nucleosome stacking and the chromatin folding at the secondary and tertiary levels

Chromatin higher-order folding is subject to regulation by posttranslational histone charge modification (for a review see (Ghoneim et al., 2021; Pepenella et al., 2014)). One of the strongest effects on global chromatin folding is imposed by arginine citrullination in the N-tails of histone H3 and H4 by the arginine deiminase PAD4, which mediates global chromatin unfolding that results in formation of neutrophil extracellular traps or NETs (Brinkmann and Zychlinsky, 2012; Wang et al., 2009). Early experiments showed that chromatin from human neutrophils underwent extensive histone proteolysis, so for experiments with PAD4-induced histone citrullination we employed chromatin purified from K562 cells. These cells are derived from immature precursors of neutrophils. They do not express the granule proteases abundant in neutrophils, but display chromatin epigenetic marks similar to that of mature neutrophils (Salzberg et al., 2017).

We isolated MNase-fragmented native nucleosome arrays and incubated them with PAD4 arginine deiminase with 2 mM CaCl_2_ (PAD4 co-enzyme) as described in the Methods. Control samples without PAD4 enzyme or without CaCl_2_ were incubated under the same conditions. The PAD4-treated and control chromatin samples were analyzed using 18% SDS-PAGE, to monitor histone citrullination by the downward mobility shifts of histone H3 and H4, due to the positive charge reduction (Wang et al., 2004). The gel staining shows a complete shift of histones H3 and H4 in PAD4-treated K562 chromatin (Fig. 5 A, lanes 2,3) but no such shift in the control sample (lane 1), indicating that the chromatin modification is fully dependent on PAD4 and not due to chromatin degradation. We confirmed this result in parallel Western blot analysis using antibodies against citrullinated histone H3 (Abcam ab5103) (Fig. 5A, lanes 5, 6 and 7), and by Triton-Acetate-Urea gels (Shechter et al., 2007) showing the upward shifts of histone H4 and H3 variants resulting from positive charge reduction (Fig 5A, lanes 8 and 9).

**Figure 5:**
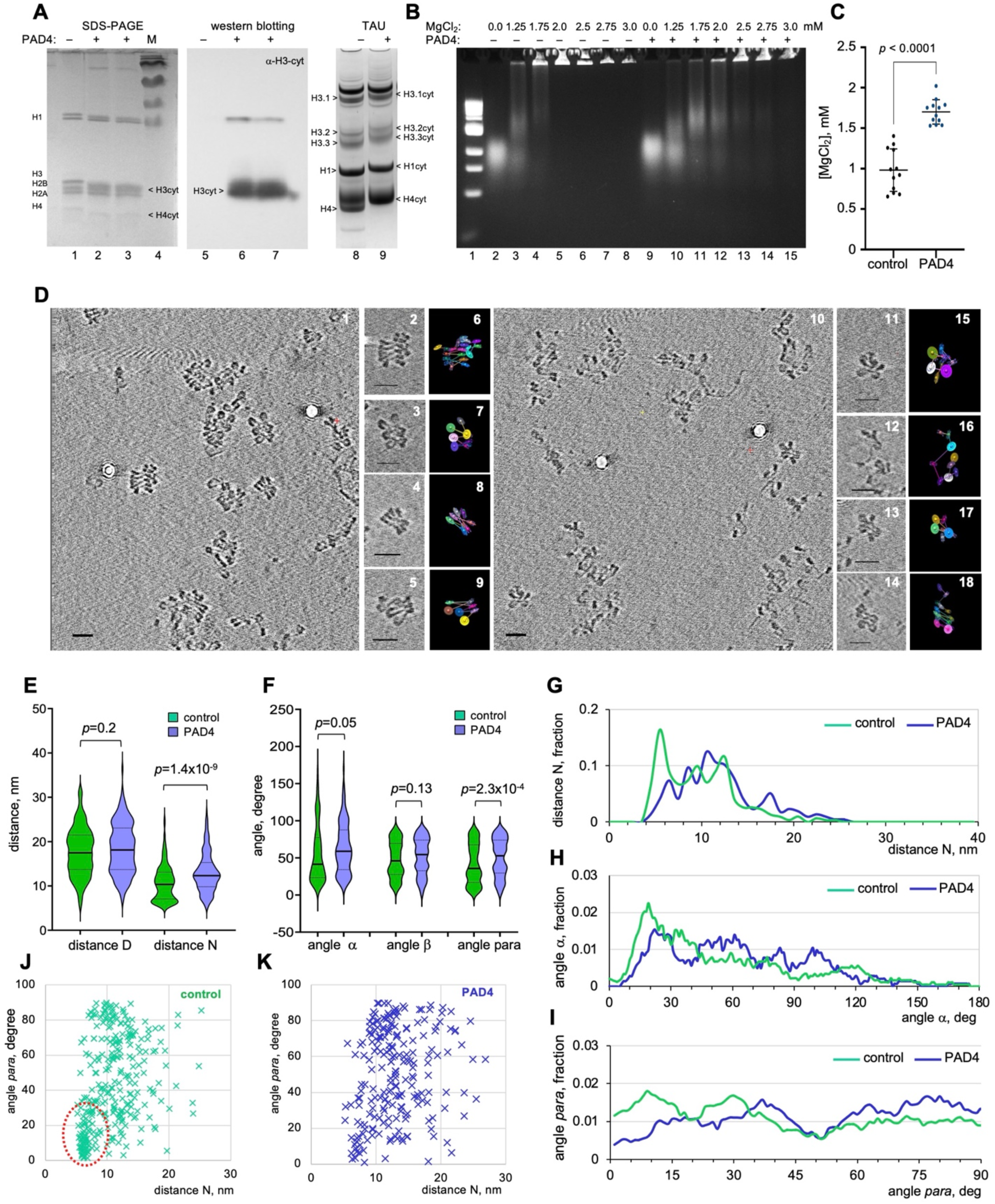
Inhibition of native chromatin higher-order folding by PAD4-dependent histone citrullination. (A) Lanes 1-3: 18% SDS-PAGE gel stained by Coomassie R250 shows histones of control (lanes 1) and PAD4-treated (lanes 2, 3) nucleosome arrays from K562 cells. Lane 4: molecular mass markers. Lanes 5-8: 8-16% gradient SDS-PAGE of histones from control (lane 5) and PAD4-treated (lanes 2, 3) nucleosome arrays followed by Western blotting detected by antibodies against histone H3 citrullinated at Arginine 2, 8, and 17. Lanes 8, 9: Triton-Acetate-Urea gel stained by Coomassie R250 shows histones extracted from control (lane 8) and PAD4-treated (lanes 9) nucleosome arrays from K562 cells. (B) DNP agarose gel showing DNA size markers (lane 1) and native electrophoresis of the control (lanes 2-8) and PAD4-treated (lanes 9-15) nucleosome arrays from K562 cells. (C) 50% self-association points determined for the control and PAD4-treated nucleosome arrays from K562 cells. (D) Panels 1-5: Cryo-ET tomogram (TS_14_2) and cropped images showing control K562 arrays vitrified at 0.75 mM Mg^2+^. Panels 6-9: the images in panels 2-5 were processed to generate CAP models (CAP#14_2_1, 14_2_2, 14_2_4, 14_1_6). Panels 10-14: Cryo-ET tomogram (TS_16_1) and cropped images showing PAD4-treated K562 arrays vitrified at 0.75 mM Mg^2+^. Panels 15-18: the images in panels 11-14 were processed to generate CAP models (CAP #16_1_8, 16_1_10, 16_1_11, 16_1_1). Scale bars: 30 nm. E-I. Violin plots (E, F) and frequency distribution profiles (G-I) of distances D, N, angle α, and angle para values calculated for control (green) and PAD4-treated (blue) nucleosome arrays vitrified at 0.75 mM Mg^2+^. Control samples: n = 209 (D), n = 302 (N), n = 162 (α), n = 302 (*para*); PAD4-treated samples: n = 162 (D), n = 215 (N), n = 130 (α), n = 125, n = 215 (para). (J, K) Two-dimensional plots showing distributions of the internucleosomal angle *para* vs. the distance N for the control (J) and PAD4-treated (K) nucleosome arrays vitrified at 0.75 mM Mg^2+^.

We then monitored the effect of PAD4 citrullination on chromatin folding by resolving crosslinked chromatin particles on a native DNP agarose gel (Fig. 5 B). Without PAD4, nucleosome arrays showed a partial self-association at 1.25 mM and complete self-association by 2.0 mM Mg^2+^ (Fig. 5 B, lanes 2-8), exactly as we observed with the untreated native chromatin (Fig. 1 C). After PAD4 treatment, we observed inhibition of chromatin folding with complete self-association well above 2 mM Mg^2+^ (Fig. 5 B, lanes 9-15). A parallel Mg^2+^-dependent precipitation assay (without glutaraldehyde crosslinking) shows the PAD4-induced change in mid-transition points (Fig. 5 C) and is consistent with the DNP gel assay.

Cryo-ET of the control and PAD4-treated native nucleosome arrays showed no substantial difference in nucleosome folding at 0 mM Mg^2+^ (suppl. Fig. S8 A, B). In the presence of 0.75 mM Mg^2+^, however, the control chromatin showed a high extent of compaction and stacking (Fig. 5 D, panel 1), which was significantly reduced in the PAD4-treated chromatin (Fig. 5 D, panel 10). At 1.0 mM Mg^2+^, the control-treated chromatin displayed clear traits of self-association (suppl. Fig. S8 E), but the PAD4-treated arrays remained dissociated (suppl. Fig. S8 F), showing that the formation of both the secondary and tertiary chromatin structures is significantly inhibited by PAD4-induced histone citrullination.

CAP modeling and quantitative stereological analysis of cryo-ET data showed that none of the recorded internucleosome distances or planes were significantly altered by PAD4 citrullination at 0 mM Mg^2+^, indicating that it did not change the nucleosome primary structure and did not cause any substantial nucleosome loss or sliding that would affect distance D (suppl. Fig. S9). At 0.75 mM Mg^2+^, however, the average distance N between the nearest neighbor nucleosomes was significantly (*p*<10^-8^) increased, from 10.7 to 13.1 nm; (Fig. 5 E) resulting in a sharp decrease in the distance N distribution peak at 6 nm, which corresponds to nucleosome disk stacking (Fig. 5 G). The distribution of distance D was not significantly affected by PAD4 treatment (Fig. 5 E), consistent with no nucleosome sliding or loss at 0.75 mM Mg^2+^.

The total average angle α value did not change significantly by PAD4 treatment (Fig. 5 F), but the angle α distribution showed a notable increase at values below 20° (Fig. 5 H). Accordingly, the lowest 20% of angles α were changed most significantly (*p*<10^-6^) while the top 20 % were not changed at all (Suppl. Fig. S10 C), consistent with histone citrullination mainly affecting stacked nucleosomes. The angle β did not show significant changes induced by histone citrullination (Suppl. Fig. S10 D, G), which is consistent with there being no reorientation between the consecutive nucleosome planes. The average angle *para* in the PAD4-treated chromatin displayed a prominent decrease at the values below 20°, and the most significant difference (*p*<10^-8^) was observed for the lowest 20% of the angle *para* populations (Fig. 5 F, I, Suppl. Fig. S10 E). Two-dimensional plotting of angles para versus the corresponding distance N values (Fig. 5 J, K) revealed the strongest change in the area below 25° angle para and at the distance N ∼ 6nm (highlighted by dashed oval), which corresponds to the tightly stacked parallel nucleosome disks. Unlike what was observed for chromatin unfolded at low salt (cf. Fig. 3 K, PAD4-induced nucleosome unfolding did not reduce the area at 60-90° angle *para* and distance N ∼ 8-12 nm, which corresponds to perpendicular nucleosome disks. This is in agreement with the lack of changes in the top quantile of angle *para* values (Suppl. Fig. S10 E). Thus, we concluded that the PAD4-induced histone citrullination was specifically inhibiting the nucleosome stacking, while having relatively little effect on the conformation of nucleosome linkers or on the spatial organization of the non-stacked nucleosomes in native chromatin.

### Reconstituted nucleosome arrays are dramatically unfolded by histone citrullination

The unfolding of native chromatin by histone citrullination notably differed from the almost complete unfolding observed for reconstituted arrays by PAD4 treatment (Wang et al., 2009). We reasoned that because histone citrullination mostly targets the nucleosome disk stacking, its effect on chromatin folding would be stronger for an array with uniform nucleosome stacking, such as that observed for clone 601-based nucleosome arrays (Garcia-Saez et al., 2018). Therefore, we constructed nucleosome arrays based on clone 601 with a NRL of 183 bp, and then reconstituted it with linker histone H1°. The 183 bp NRL was chosen as a representative of the 10n+5 class of nucleosome linkers (close to 172, 183, and 193 bp), which is typical of natural chromatin (Grigoryev, 2012).

The 12-mer, 183 bp NRL nucleosome core arrays were reconstituted, isolated, and characterized as previously described (Bass et al., 2019; Grigoryev et al., 2016). Arrays were treated with PAD4 arginine deiminase (Sigma SRP0329) and compared to untreaded samples by 18% SDS-PAGE, to confirm a complete shift of histones H3 and H4 (Fig. 6 A, lanes 2,4). No such shift was observed in the control samples (lanes 1, 3). PAD4-treated and control nucleosome arrays were then mixed with recombinant linker histone variant H1° (New England BioLabs; catalog no. M2501S) as described before (Grigoryev et al., 2016), at a ratio of 0.7 molecules per nucleosome, which is close to the natural stoichiometry of linker histone H1 in most proliferating cells (Woodcock et al., 2006). We reconstituted linker histone H1 with the already citrullinated nucleosome arrays because PAD4 partial citrullinates histone H1 (Fig 5A), and we sought only to monitor the effect of core histone citrullination.

**Figure 6:**
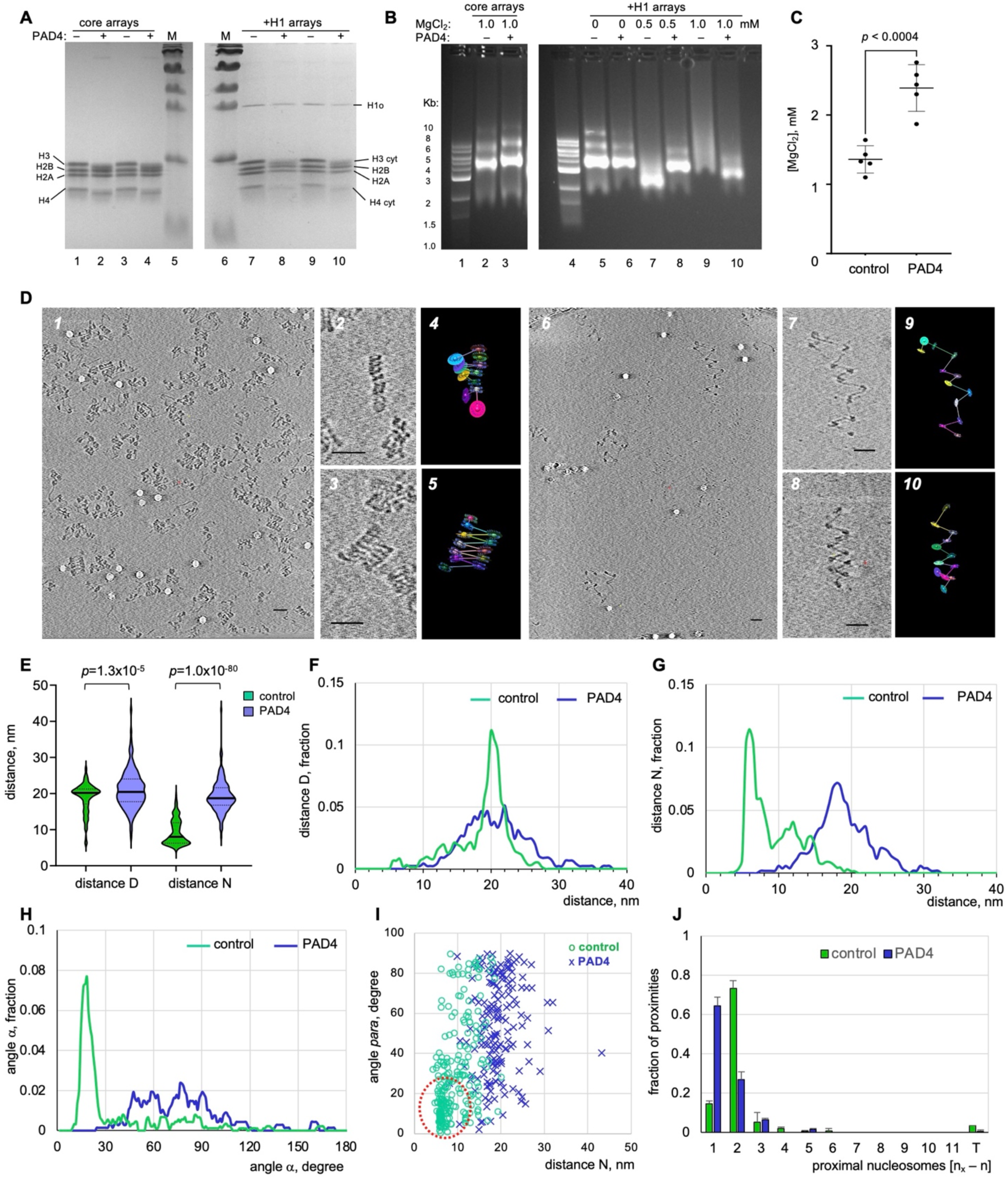
Nucleosome chain folding in clone 601-based reconstituted nucleosome arrays and its inhibition by histone citrullination. (A) Lanes 1-5: 18% SDS-PAGE gel stained by Coomassie R250 shows histones of control (lanes 1, 3) and PAD4-treated (lanes 2, 4) reconstituted nucleosome core arrays. M – m.w. markers (lane 5). Lanes 1-5: 18% SDS-PAGE gel stained by Coomassie R250 shows histones of control (lanes 7, 9) and PAD4-treated (lanes 8, 10) reconstituted nucleosome arrays additionally reconstituted with linker histone H1° after PAD4 treatment. (B) DNP agarose gels showing DNA size markers (lanes 1, 4) and native electrophoresis of the control (lane 2) and PAD4-treated (lanes 3) reconstituted nucleosome core arrays crosslinked with glutaraldehyde at 1 mM Mg^2+^ and control (lanes 5, 7, 9) and PAD4-treated (lanes 6, 8, 10) reconstituted 183×12+H1 arrays after PAD4 treatment crosslinked at 0 to 1 mM Mg^2+^ as indicated. (C) 50% self-association points determined for the control and PAD4-treated nucleosome core arrays and the 183×12+H1 arrays. (D) Panels 1: Cryo-ET tomogram (TS_17_6) showing control reconstituted 183×12+H1 nucleosome arrays vitrified at 0.75 mM Mg^2+^. Panels 2-5: the cropped images (2, 3) were processed to generate CAP models CAP #17_6_1 (4), 19_2_1 (5), Panels 6-8: Cryo-ET tomogram (TS_18_3) showing PAD4-treated reconstituted 183×12+H1 nucleosome arrays vitrified at 0.75 mM Mg^2+^. Panels 7-10: the cropped images (7, 8) were processed to generate CAP models (CAP#18_1_1 (9), 18_3-2 (10). Scale bars: 30 nm. (E) Violin plots showing distribution of internucleosomal distances D and N calculated for control (green) and PAD4-treated (blue) reconstituted +H1 nucleosome arrays vitrified at 0.75 mM Mg^2+^. (F–H) Frequency distribution profiles of distances D (panel F), N (G) and angles α (H) measured for arrays of control (green curves) and PAD4-treated (blue curves) reconstituted 183×12+H1 nucleosome arrays vitrified at 0.75 mM Mg^2+^. Control samples: n = 204 (D), n = 230 (N), n = 197 (α); PAD4-treated samples: n = 150 (D), n = 158 (N) n = 135 (α). (I) Two-dimensional plot showing distribution of the internucleosomal angle *para* vs. the distance N for the control (green) and PAD4-treated (blue) reconstituted +H1 nucleosome arrays vitrified at 0.75 mM Mg^2+^. (J) Distribution of pairwise nucleosome proximities determined for the control (green columns) and PAD4-treated (blue columns) reconstituted +H1 nucleosome arrays vitrified at 0.75 mM Mg^2+^. Control samples: n = 237; PAD4-treated samples: n = 165.

We monitored the effect of PAD4 citrullination on the reconstituted nucleosome arrays by resolving crosslinked arrays DNP agarose gels (Fig. 6 B). Without linker histone H1, PAD4 treatment caused only a slight upward shift of the 183×12 core arrays, suggesting partial chromatin unfolding due to histone citrullination (Fig 5 B, lanes 2, 3). We observed a strong downward shift in H1-reconstitued arrays at 0.5 mM Mg^2+^ (lanes 7) and a prominent band smearing at 1 mM Mg^2+^ (lane 9), while the PAD4-treated arrays showed almost complete inhibition of both downward shift (lane 8) and band smearing (lane 10). In parallel, we conducted Mg^2+^-dependent self-association assays showing the PAD4-induced change in the mid-transition points of the 183×12+H1 arrays, which is fully consistent with the DNP gels (Fig 6 C). Thus, we concluded that PAD4-mediated histone citrullination had a profound inhibitory effect on the secondary and tertiary chromatin structures formed with reconstituted arrays in the presence of linker histone, and that physiological Mg^2+^ concentrations have a very similar effect to that observed in native chromatin.

Cryo-ET showed that the control 183×12+H1 nucleosome arrays exhibited much more abundant and uniform nucleosome stacking at 0.5 mM Mg^2+^ (Fig. 6 D, panels 1-3) than was observed with native chromatin above. Strikingly, despite the presence of 0.5 mM Mg^2+^, reconstituted arrays underwent complete unfolding upon treatment with PAD4, showing extended zigzag arrays of nucleosomes (Fig. 6 D, panels 6-8). This observation is in full agreement with the solution assays, showing that the formation of both the secondary and tertiary chromatin structures is abrogated by PAD4-induced histone citrullination at physiological Mg^2+^ concentration.

CAP modeling and stereological analysis of control and PAD4-treated 183×12+H1 arrays at 0.5 mM Mg^2+^ showed that the average distance N between the nearest neighbor nucleosomes increased more than 2-fold (Fig. 6 E), from 9.3 to 19.3 nm (*p*<10^-80^) (i.e., dramatically stronger than in the PAD4-treated native chromatin, cf. Fig. 5 E). This leads to a complete disappearance of the distance N distribution peak at 6 nm (Fig. 6 G). The average distance D was not as strongly affected by PAD4 treatment (Fig. 6 E), however, the distribution showed a striking change between the sharp peak centered at 21 nm for control chromatin to a wide distribution in the PAD4-teated chromatin (Fig. 6 F). These results likely reflect the wide conformational flexibility of linker DNA in the unfolded state.

Similarly, the angle α distribution showed a dramatic loss of its major peak at ∼18° and the appearance of a very wide distribution in the PAD4-treated arrays (Fig. 6 H). The distribution of angle β showed rather modest changes resulting from histone citrullination, though the reduced angle β values below 15° and increased values above 70° were notable (Suppl. Fig. S11 B, C).

The average angle *para* in the PAD4-treated chromatin displayed highly significant differences (*p*<10^-19^) and a prominent decrease at the values below 20° (Suppl. Fig. S11 B, D). A two-dimensional plot of angles *para* versus the corresponding distance N values (Fig. 6 I) reveals an obvious change in the area below 30° angle *para* and at the distance N ∼ 6nm (dashed oval). Accordingly, the peak of nucleosome proximities at i±2, which corresponds to the predominately zigzag folding, was dramatically reduced by PAD4 treatment in parallel to an equally strong increase in i±1, which corresponds to an extended nucleosome chain (Fig. 6 J). Thus, the reconstituted nucleosome arrays showed a dramatic unfolding by PAD4-mediated histone citrullination, exceeding that in the native chromatin and consistent with complete disruption of the overall nucleosome disk stacking in the reconstituted clone 601-based arrays.

### Nanoscale spatial analysis reveals fundamental topological differences between the native and reconstituted nucleosome arrays folding

Since most previous chromatin structural studies have focused separately on either native (Horowitz et al., 1994; Kizilyaprak et al., 2010; Scheffer et al., 2011) or reconstituted chromatin (Dorigo et al., 2004; Garcia-Saez et al., 2018; Routh et al., 2008; Schalch et al., 2005; Song et al., 2014), here we compared the overall degree of compaction and the intrinsic nucleosome folding paths in the native and reconstituted chromatin samples by Cryo-ET, under the same conditions (0.75 mM Mg^2+^). Comparison of the average distance between the consecutive nucleosomes, D did not show significant changes between the native and reconstituted chromatin (Fig. 7 A), however, the distribution of distances showed a striking contrast between the sharp peak at 21 nm in the 183×12 reconstitutes and several broad peaks in the native K562 chromatin (Fig. 7 C). This striking difference is likely due to the wide variability in linker DNA length in native K562 chromatin, as well as the conformational uniformity in the DNA linkers observed for the reconstituted nucleosome arrays by Cryo-ET (Fig. 6 D, panels 1-3). The average distances N between the proximal nucleosomes (Fig. 7 A) showed a modest increase, which is consistent with the increase observed in the broad distribution peak at ∼ 11 nm in the native chromatin (Fig. 7 D). As we pointed above, this peak in N corresponds to the nucleosomes contacting each other at a perpendicular orientation (Fig. 3 L, M). Such nucleosomes are very rare in the reconstituted nucleosome arrays by Cryo-ET (Fig. 6 D, panels 1-3).

**Figure 7:**
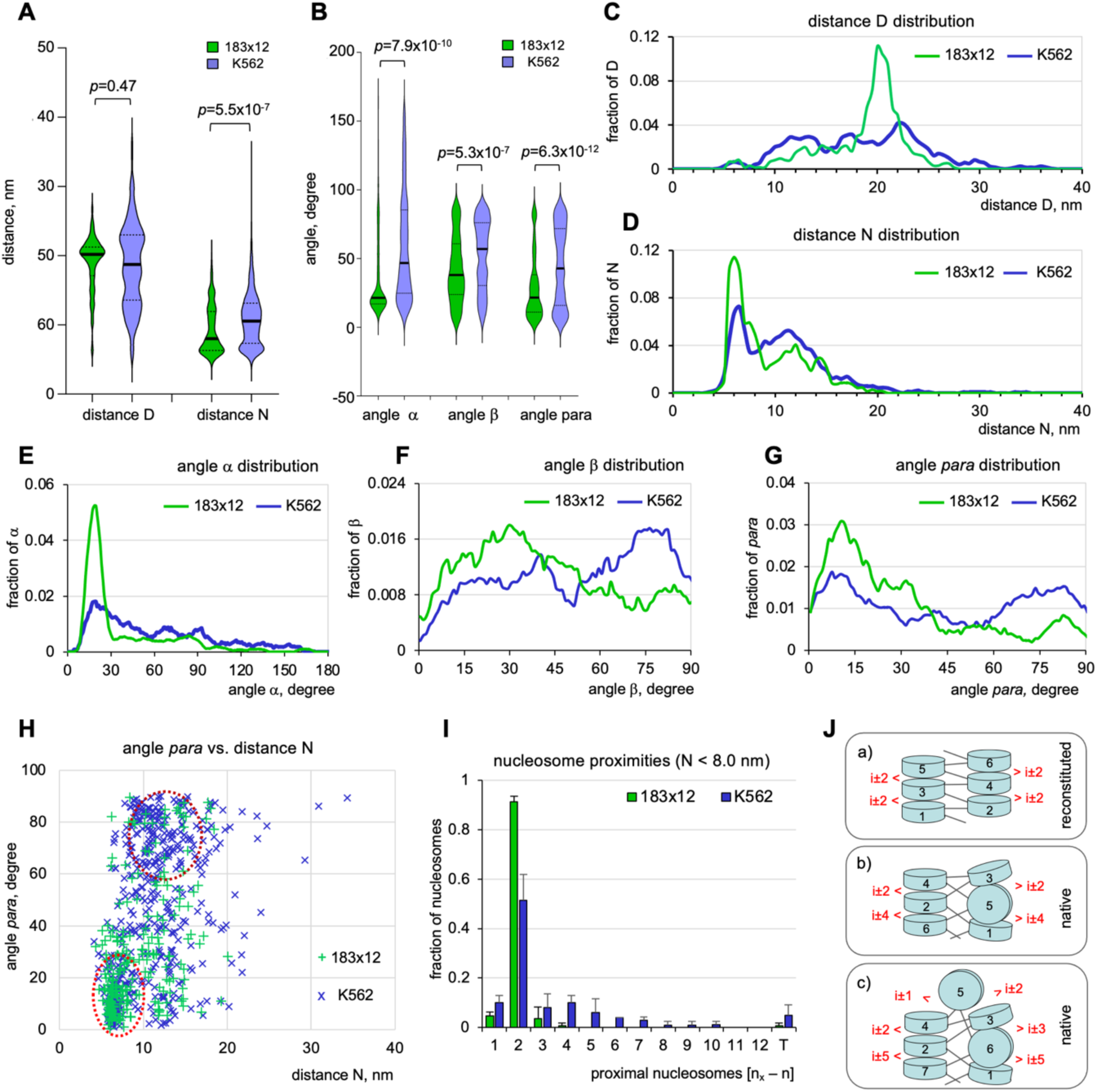
Comparative stereological analysis of the native and reconstituted chromatin folding. (A, B) Violin plots comparing distributions of distances D and N (A) and angles α, β, and *para* (B) determined for the 183×12+H1 nucleosome arrays (green) and native K562 chromatin (blue) vitrified at 0.75 mM Mg^2+^. (C-G) Frequency distribution profiles comparing distances D (panel C), N (D) and angles α (E), β (F), and *para* (G) measured for the 183×12+H1 nucleosome arrays (green) and native K562 chromatin (blue) vitrified at 0.75 mM Mg^2+^. (H) Two-dimensional plot showing distribution of the internucleosomal angle *para* vs. the distance N for the 183×12+H1 nucleosome arrays (green) and native K562 chromatin (blue) vitrified at 0.75 mM Mg^2+^. (I) Distribution of pairwise nucleosome proximities determined for the 183×12+H1 nucleosome arrays (green) and native K562 chromatin (blue) vitrified at 0.75 mM Mg^2+^. (J) Cartoon models illustrating the paths of nucleosome chain folding in the reconstituted arrays (a) and two types of native nucleosome arrays (b, c).

The axial angle α and plane angles β and *para* were all significantly different between the native and reconstituted chromatin (Fig. 7 B). The angle α distribution showed a maximum at ∼18° in both samples, but had a much higher and narrower peak in the reconstituted chromatin (Fig. 7 E). The values of angle α between 60° and 100° were more prominent in the native chromatin. Notably, this range of angles dominates in the unfolded chromatin (Fig. 3 G), indicating that this component of angle α distribution reflects the presence of unfolded linker DNA in the Mg^2+^-condensed native chromatin, in contrast to reconstituted arrays. This incomplete folding may originate from partial unfolding of individual nucleosome cores in K562 chromatin such as those with angle α more than 90° that can be seen, e.g., for the two left nucleosomes in Fig. 3 A, B. The angle β was rather widely distributed in both samples. The largest difference with native chromatin was manifested by the notably diminished peak at 60-90° in reconstituted arrays (Fig. 7 F), which represents consecutive nucleosomes with planes oriented perpendicularly (e.g., nucleosome #4 and #5 on Fig. 7 J “native” cartoons b and c).

The angle *para* in the native chromatin also displayed a prominent decrease at the values between 60° and 100° in the reconstituted vs. native chromatin (Fig. 7 G). The two-dimensional plotting of angles *para* vs. the corresponding distance N values (Fig. 7 H) shows that the clustered area below 30° and at the distance ∼ 6nm (bottom dashed oval) is very prominent in both samples. The other clustered area at 60-90° angle *para* and distance N ∼ 8-12 nm (top dashed oval), corresponding to perpendicular nucleosome disks, is largely absent in the reconstituted nucleosome arrays. This difference is fully consistent with cryo-ET data showing that proximal nucleosomes with planes oriented perpendicularly (e.g. nucleosome #3 and #5 on Fig. 7 J “native” cartoons b and c) are absent from the reconstituted nucleosome arrays (Fig 7 J, cartoon a).

When we compared nucleosome proximities in the Mg^2+^-condensed native chromatin (Fig. 3 J) and reconstituted arrays (Fig. 6 J), we were intrigued by the presence of prominent proximities at i±3, i±4 and i±5 in the native chromatin, which were virtually absent in the reconstituted samples. This difference could be the product of less tightly folded native chromatin or a fundamental difference in the nucleosome chain topology. To distinguish between the two possibilities, we subtracted all unfolded nucleosomes (with distance N > 8.0 nm) from both data sets, and the observed trend was fully retained for these closely juxtaposed nucleosomes (Fig. 7 I). Upon revisiting the native nucleosome structure models and datasets (Suppl. Fig. S4 B and Suppl. Table S2), we found that the i±3, i±4 and i±5 interactions originated from a special folding of the native nucleosome arrays that deviated from the standard zigzag path and, after making a U-turn, formed inverse zigzag folds with crisscrossed linker DNA (e.g. Fig. 3 D, M). Depending on the even or odd number of nucleosomes making the U-turn, such structures would either display i±4 (Fig. 7 J, “native” cartoon b) or i±1, i±3 and i±5 (Fig. 7 J, “native” cartoon c) proximities, in addition to the most dominant i±2. In contrast, the reconstituted arrays, which form a classic two-start zigzag fiber, are expected to display only i±2 proximities (Fig. 7 J, “reconstituted” cartoon a) consistent with Cryo-ET observations (Fig. 6 D). Thus, nanoscale spatial analysis of native chromatin folding revealed three novel features specific to the condensed native chromatin: 1) the discontinuous asymmetric nucleosome stacking; 2) the alternating parallel and perpendicular orientations of the juxtaposed nucleosome disks; and 3) the inverse zig zag topology of nucleosome chains that, despite displaying prominent i±2 proximities, fold into discontinuous nanoparticles rather than continuous zig zag fibers.

## DISCUSSION

Our Cryo-ET analysis combined with stereological nanoscale modeling reveals, for the first time, the regressive zigzag path of the nucleosome chain in native condensed chromatin. These findings help to reconcile long-standing controversies regarding the difference in chromatin folding *in vitro* and in the interphase cell, and provide novel mechanistic insights into the cooperation of distinct structural motifs in the higher-order folding of the nucleosome chain. As it was recently observed by live cell imaging, the nucleosomes may fold into relatively small (4-8 nucleosome) clutches or nanodomains instead of fibers (Kieffer-Kwon et al., 2017; Nozaki et al., 2017; Ricci et al., 2015). Whether these nanodomains have any particular structure above the primary nucleosome chain remained unknown. Here, based on the results of Cryo-ET, we propose a new general model for the nucleosome chain folding into a discontinuous secondary structure comprised of stacks of closely juxtaposed nucleosome disks interrupted by nucleosomes oriented perpendicularly to their neighbors, thus breaking the continuous nucleosome stacking (Fig. 8).

**Figure 8:**
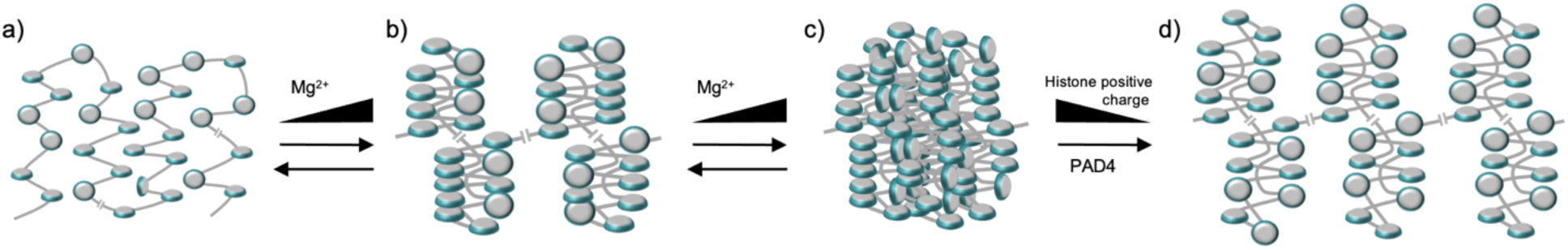
Schematic model for nucleosome chain folding into the secondary and tertiary higher-order structures mediated by nucleosome stacking and reversed by histone positive charge reduction. a) primary structure: unfolded nucleosome chains; b) secondary structure: nanoparticles mediated by nucleosome stacking in-cis; c) tertiary structure: the nanoparticles are joined together by nucleosome stacking *in-cis* and *in-trans*; d) complete unfolding of the tertiary structure and partial unfolding of the secondary structure by histone modifications (such as histone citrullination or acetylation) reducing the net positive charge of the histones.

We found that upon the Mg^2+^-induced compaction the axial angle α and the internucleosomal angle *para* were reduced dramatically in both native (Fig. 3 G, I) and reconstituted chromatin (Fig. 6, H,I). These observations are consistent with previous models where contraction of angle α is mediated by linker histone H1 and electrostatic stabilization of the juxtaposed linker DNA by monovalent or divalent cations (Grigoryev et al., 2009; Grigoryev et al., 2016). Unlike the longitudinal accordion-like folding predicted by the modeling and observed in reconstituted arrays (Fig. 6), we found that the nucleosome chain path in condensed chromatin makes frequent fold-backs and is engaged in a regressive zig zag folding. In this model, the stacked nucleosomes disks alternate with those that maintain open nucleosome surfaces (Fig. 8b), resulting in a wide variation in the stacking rate within individual nucleosome arrays (Suppl. Fig. S4 B). This cooperative nucleosome stacking clearly presents the major energetic barrier for chromatin unfolding and is strongly dependent on the linker DNA length (Brouwer et al., 2021; Dombrowski et al., 2022). The nucleosome surfaces including the unique acidic patch that become closed by the nucleosome stacking are the sites of interaction with multiple chromatin regulatory and architectural factors (Skrajna et al., 2020). Whether nucleosome arrays with the maximal or minimal percentage of stacking originate from repressed and active chromatin domains and/or regions with specific nucleosome positioning requires further investigation.

The efficiency of chromatin fiber folding in reconstituted arrays was previously shown, by us, to depend on DNA linker length and the associated angle β (Correll et al., 2012). In reconstituted arrays, however, the uniform nucleosome linker lengths were generated to impose uniform angles β (either positive or negative) between the nucleosome planes. This helped to mediate a progressive increase in the nucleosome chain length upon the increase in the nucleosome cores. If the linker DNA generates both positive and negative angles β, then the nucleosome path will accordingly reverse direction and fold back upon itself. In this work we were only able to record the absolute values of angle β, so its role in generating the regressive zig zag path of the nucleosome chain remains an open question. This intriguing possibility may be tested by engineering new nucleosome arrays with variable linker DNA lengths and angles β designed to reverse the nucleosome chain path.

Using *in situ* EMANIC we previously showed that reconstituted nucleosome chains, as well as chromatin from terminally differentiated cells, exhibited uniform i±2 interactions, which was consistent with the two-start zig zag fiber. In human interphase cells the nucleosome chain was shown mostly to make i±2 interactions with additional i±1, i±3, i±4, and i±5. This pattern was interpreted as interphase chromatin folding into a continuous but heteromorphic zig zag containing different linker DNA conformations within the same fiber. These EMANIC data are consistent with genome-wide studies of nucleosome proximities by radiation-induced spatially correlated cleavage of DNA (Risca et al., 2017), or interactions by Micro-C (Hsieh et al., 2020; Krietenstein et al., 2020), which all showed a distinct pattern of i±2 nucleosome proximities in addition to several other near-neighbor interactions. In all these cases, however, the nucleosome interactome was concluded to be evidence for the two-start zig zag folding while the nature of its deviation from the pure zig zag models remained unclear.

Here, the direct imaging of nucleosome chain folding by Cryo-ET showed a pattern of nucleosome proximities, in native chromatin, that are remarkably similar to that of *in situ* EMANIC (cf. Fig. 3 L Suppl. Fig. S6 G), as well as the other interaction mapping studies cited above. Clearly, however, the path deviates from that of reconstituted nucleosome arrays which have uniform i±2 proximities (Fig. 7 I). Our data are consistent with higher flexibility along the chromatin zig zag path, which negates the concept of a continuous 30-nm fiber within interphase nuclei (Fig. 8b). Furthermore, our Cryo-ET imaging of discontinuous nucleosome stacks is consistent with nanodomains observed by super-resolution microscopy (Fang et al., 2018; Kieffer-Kwon et al., 2017; Ricci et al., 2015) and with the short patches of nucleosome stacks, hubs, and loops observed by EM tomography of chromatin in situ (Ou et al., 2017). Our new findings resolve the path of linker DNA and individual nucleosome interactions in condensed chromatin and provide experimental grounds for developing new 3D nucleosome chain models. They will also inform dynamic simulations of internucleosomal interactions for better understanding of the genome-wide nucleosome interactome, as well as aid in designing small molecules targeting nucleosome interactions in a physiological setting.

Previous TEM studies of Mg^2+^-condensed reconstituted nucleosome chains have revealed large globular structures, but they did not reveal any distinct structural elements, in particular, such as 30-nm fibers (Maeshima et al., 2016). More recently, the mechanism of Mg^2+^-dependent condensation and self-association of nucleosome arrays has been described as a liquid-liquid phase separation (LLPS) where the condensed nucleosome arrays are expected to reside in a fluid unstructured state (Gibson et al., 2019). With native chromatin extensively condensed at Mg^2+^ concentration higher than 1 mM, our TEM experiments also failed to detect any internal folding of the nucleosomes (Fig. 1 D). Remarkably, however, the nucleosome condensates formed in the presence of either 1.25 or 2 mM Mg^2+^ appeared to flatten rapidly in the process of vitrification, which is consistent with a highly fluid nature of the nucleosome condensates (Gibson et al., 2019), and they were readily observed by Cryo-ET (Fig. 2).

Despite this apparent fluidity, Cryo-ET showed patches of stacked nucleosomes, demonstrating that the secondary structure motifs are not completely erased during tertiary chromatin folding. This raises the interesting possibility that the cooperative nucleosome self-association forms nanodomains that maintain their compact state within the condensed “liquid droplets”, like those formed during mitosis (Schneider et al., 2021), and thus retain epigenetic “memory” of their repressed state. Within the self-associated nucleosome condensates, we observed multiple internucleosomal trans-contacts mediated by anti-parallel stacking of nucleosome disks. Such contacts are consistent with the nucleosome interdigitation hypothesis previously proposed as a mechanism of nucleosome self-association in condensed heterochromatin (Grigoryev, 2004) and metaphase chromosomes (Chicano et al., 2019).

Unlike these idealized models, however, in nucleosome condensates observed by Cryo-ET, antiparallel nucleosome stacking did not show a tendency to dominate over the stacks of parallel nucleosomes, and they formed a minority of all visible stacking interactions (Fig. 4). We suggest that in the process of Mg^2+^-mediated formation of the tertiary chromatin structure, the nucleosome surfaces that remain open in the secondary structure (Fig. 8b) would interact *in-trans* and form contacts between the antiparallel nucleosome surfaces leading to the bulky nucleosome condensates (Fig. 8c). Apparently newly formed parallel nucleosome stacking *in-trans* requires a higher Mg^2+^ concentration than the *in-cis* stacking. It had been noticed before that a number of nucleosome core crystal lattices display anti-parallel “head-to-tail” nucleosome stacking consistent with *in-trans* nucleosome interactions within the self-associated nucleosome condensates (Grigoryev, 2004; Korolev et al., 2018). In particular, such stacking was observed by X-ray crystallography of nucleosomes containing histone variant H3.Y, which also feature a partial unfolding of nucleosome ends and reduced binding to linker histone H1 (Kujirai et al., 2016). It remains to be shown whether the nucleosomes that resist *in-cis* stacking in the secondary structure but are engaged in *in-trans* stacking in the tertiary structure bear any specific histone variants or other epigenetic marks regulating the process of global chromatin condensation.

The Mg^2+^-induced native chromatin folding and formation of the secondary structure at 0.5 – 0.75 mM Mg^2+^ (Fig. 2) is remarkably close to the physiological concentration of free Mg^2+^, estimated at ∼0.6 mM in the interphase cell (Maeshima et al., 2018). Further increase in Mg^2+^ promotes metaphase chromosome condensation *in vivo* (Dwiranti et al., 2019; Maeshima et al., 2018) by a process very similar to the Mg^2+^-dependent self-association *in vitro* (Maeshima et al., 2016). We thus assume that the Mg^2+^- induced structural transitions of native chromatin observed by Cryo-ET correctly reproduce those observed *in vivo* at similar Mg^2+^-concentrations. Previously we have simulated two interdependent effects of Mg^2+^ on chromatin folding: 1) the electrostatic screening of phosphate groups in DNA facilitating nucleosome core juxtaposition, and 2) promoting partial linker DNA bending, which helps to resolve spatial clashes in the chromatin fiber (Grigoryev et al., 2009). Both of these model predictions are consistent with the new Cryo-ET studies. Going beyond the model predictions and experiments, however, here we observed a surprisingly higher heterogeneity of nucleosome stacking in native chromatin (Fig. 3, Suppl. Fig. S4 B).

In addition to the uniform electrostatic screening of negative charges in DNA, Mg^2+^ has been shown to alter chromatin folding in a sequence-specific manner by promoting selective self-assembly of nucleosomes bearing identical DNA sequences (Nishikawa and Ohyama, 2013). This, still enigmatic, effect of Mg^2+^ on chromatin folding has been proposed to be gene-specific and is thought to facilitate nucleosome stacking and condensation of heterochromatin enriched by highly repetitive satellite DNA (Ohyama, 2019). Further work on Mg^2+^-induced compaction of fractionated native chromatin, followed by Cryo-ET and DNA sequence analysis, should reveal the nature of these genomic regions and chromatin states enriched or depleted in the extent of nucleosome stacking.

The unstructured N-terminal tails of histones H3 and H4 have been shown to mediate internucleosomal interactions (both *in-cis* and *in-trans*), contributing to the higher-order chromatin folding. Because of this they are targets for chromatin structure regulation by posttranslational histone charge modifications (for review see (Ghoneim et al., 2021; Pepenella et al., 2014)). One of the strongest effects on global chromatin folding is imposed by arginine citrullination in histone H3 and H4 N-tails by the arginine deiminase PAD4. Activation of PAD4 in neutrophil granulocytes causes massive chromatin unfolding during NETosis (Brinkmann and Zychlinsky, 2012; Wang et al., 2009). Here, using Cryo-ET we found that PAD4 significantly inhibits the Mg^2+^ -dependent native chromatin folding at both the secondary and the tertiary levels, by disrupting the nucleosome stacking interactions. Accordingly, the extent of secondary structure unfolding was reduced in native chromatin, which contained only about 30% stacked nucleosomes compared to reconstituted arrays, which show almost complete nucleosome stacking (cf. Figs. 5 and 6). The chromatin in mature human granulocyte nuclei is highly condensed and accumulates abundant tertiary structures (Xu et al., 2021) before acquiring the ability to release NETs in the process of granulocyte maturation (Lukasova et al., 2013). We propose that a major effect of PAD4-induced citrullination is to reduce the tertiary chromatin structure by modifying the histone H3 and H4 tails, thus inhibiting nucleosome stacking interactions at the physiological Mg^2+^ concentration. This structural transition, though insufficient by itself to unfold the chromatin into NETs many times larger than the nuclear diameter, may trigger NETosis by unfolding the nucleosome condensates, and opening them to additional factors, such as calpain protease (Gosswein et al., 2019), which would further promote chromatin decondensation by digesting histone tails and other nuclear architectural proteins that lead to complete nuclear rupture (Thiam et al., 2020).

The fact that PAD4-mediated NETosis, in addition to its benefits for wound healing (Brinkmann and Zychlinsky, 2012), has strong adversary effects contributing to deep vein thrombosis (Martinod et al., 2013), cancer metastasis (Yang et al., 2020), and COVID-19 pathology (Barnes et al., 2020; Veras et al., 2020), makes targeting the molecular interactions leading to the release of NETs an important goal in biomedicine. One of the approaches to regulate or prevent the PAD4-mediated NETosis would be to genetically alter histone modifications and other chromatin architectural factors that potentiate granulocytes to NETosis upon maturation. Another approach would be to find a way of preventing chromatin unfolding by increasing free Mg^2+^ in neutrophils since a very small increase may efficiently preserve chromatin in the condensed state (Fig. 2 B). Based on these results, cryo-ET combined with nanoscale stereological modeling should be an immense tool for detecting chromatin structural transitions underlying chromatin functions, and a valuable resource for further analysis of the convoluted chromatin path observed by EM or super resolution microscopy *in situ*.

## ACKNOWLEDGMENTS

We are thankful to J. Sloppy and H. Chen for technical assistance with electron microscopy at the Penn State Hershey Imaging Facility. We thank J. Buckwalter and L. Gautam (Penn State Hershey) for research assistance on earlier aspects of this project. Support for this work was provided to S. Grigoryev by NSF grant 1911940 and by Pennsylvania Department of Health using Tobacco Settlement Funds Grant 4100072562. The Department specifically disclaims responsibility for any analyses, interpretations, or conclusions.

## METHODS

### Cells and tissue isolation

Human K562 cells (ATCC CCCL-243) were grown at 37°C and 5% CO_2_ in RPMI 1640 medium +GlutaMAX^TM^(Gibco 61870-036) supplemented with 10% FBS (HyClone SH30071) and Penicillin/Streptomycin in 600 mL (150 cm^2^) tissue culture flasks with passing by dilution into new media every 72 hours. For preparative chromatin isolation, cells were grown in suspension until reaching density of 0.8-1.2 x 10^6^ cells/mL to a final volume of 400 mL. The detached K562 cells were collected and twice washed by spinning down and resuspending in PBS. To block mitosis in the G1 and S-phases, K562 cells were treated with 0.5 mM mimosine or 0.5 mM hydroxyurea for 24 hr prior to nuclei isolation.

Human HeLa S3 Cells (ATCC # CCL-2.2) were grown at +37°C and 5% CO_2_ in RPMI 1640 medium +GlutaMAX^TM^ (Gibco 61870-036) and 10% FBS (HyClone SH30071.03) in 10 cm^2^ tissue culture plates with 10 mL of medium and passing by trypsinization and dilution into new media every 3-4 days. For preparative chromatin isolation, cells were grown to ca. 90% confluence in a final volume of 100 mL (10 plates), gently washed by pipetting, and detached by cell scraping in RSB buffer (10 mM NaCl, 3 mM MgCl_2_, 10 mM HEPES, pH=7.5) containing 0.5% Nonidet P-40 in RSB, 1 mM PMSF, and protease inhibitor cocktail (Sigma P8849).

Wild-type C57Bl/6J mice were purchased from The Jackson Laboratory (strain no. 000664). All breeding and experimental procedures were carried out in accordance with the guidelines set by the National Institutes of Health’s Office of Laboratory Animal Welfare and the National Science Foundation. All experiments were approved by the Penn State University College of Medicine (protocol no. 00929). Mouse retina was collected for postnatal day 21 mice as previously described (Popova et al., 2013).

### Isolation of nuclei and native chromatin

K562 and HeLa cell nuclei were isolated by resuspending the cells in 30 mL RSB buffer containing 0.5% Nonidet P-40, 1 mM PMSF, and protease inhibitor cocktail (Sigma P8849) at 4 °C as described (Grigoryev et al., 2016). The cell suspensions were homogenized by 30 strokes of pestle B in a Dounce homogenizer over 30 min on ice. Nuclei were centrifuged for 5 min at 4000 rpm and 4oC in an Eppendorf 5810 R centrifuge with an Eppendorf A-4-62 swinging bucket rotor, and the nuclear pellets were resuspended in 10 mL RSB plus 0.5 mM PMSF. Average yield for K562 and HeLa nuclei ∼1 mg/mL and ∼0.5-0.7 mg/mL of DNA, respectively, determined spectrophotometrically. The isolated nuclei were warmed at 37oC for 5 min and treated with micrococcal nuclease (MNase) (Roche 10107921001) and 2 mM CaCl_2_ (RICCA R1760000). MNase dilutions were prepared fresh before each digestion at 25 μg/mL in 1X bovine serum albumin (BSA) (New England Biolabs B9001S).

Prepared MNase/BSA solution was then added to nuclei suspension at a dilution of 1:1000 and incubated for 60 minutes at 37oC in a Belly Dancer Orbital Shaker (IBI BDRAA115S) at 40-45 rpm to prepare chromatin fragments with a maximal median DNA size of ∼2400 bp corresponding to 12 nucleosomes. The size of the nuclease fragments was determined by agarose DNA electrophoresis after DNA deproteinization by 1% SDS and 0.5 mg/ml proteinase K. The MNase digestion was stopped by adding 5 mM EDTA, rapidly cooling on ice, and spinning down for 5 min. at 10,000 rpm and 4oC in an Eppendorf 5810 R centrifuge with F-34-6-38 rotor. The nuclear pellet was resuspended in TE buffer (10 mM Tris, 1 mM EDTA, pH=8.0) and incubated at 4oC for 24-48 hr with mild shaking at the start and end of incubation. The nuclear material was then centrifuged for 5 min. in the same manner at 10,000 rpm and 4oC. The supernatant (S2) containing soluble native chromatin was collected and the concentration of soluble DNA was monitored spectrophotometrically.

The released soluble chromatin S2 was concentrated on Amicon ®Ultra-15 mL centrifugal filters (Millipore UFC9050) to 0.35-0.5 mL with the final concentration ranging from 4.0-8.0 mg/mL. Concentrated chromatin samples were loaded on 11 mL gradients of 5-25% sucrose in TE buffer and centrifuged in a preparative ultracentrifuge with a SW41 Ti rotor (Beckman) for 11 hrs. at 35,000 rpm and 4oC. Aliquots of the sucrose gradient fractions were treated with 0.5 mg/mL Proteinase K in 1% SDS for 2 hr. at 55oC, and DNA analyzed by electrophoresis in 1% agarose in TAE buffer as before (Grigoryev et al., 2016). Fractions enriched in particles with DNA sizes corresponding to 8-16 nucleosomes were collected, dialyzed for 48 hr. in a 1:250 ratio against HNE buffer (5 mM NaCl, 0.1 mM EDTA, 10 mM HEPES-NaOH, pH=7.5) in Spectra/PorTM 1 RC Dialysis Membrane Tubing (SpectrumTM 132650) and concentrated in Amicon ®Ultra-2 mL centrifugal filters (Millipore UFC205024).

Isolation of nuclei and native soluble chromatin from mouse retina tissue was carried out as previously described (Popova et al., 2013).

### Reconstitution and biochemical analysis of the nucleosome arrays

Procedures for construction of DNA templates (183×12) for multimeric oligonucleosome arrays, their reconstitution with core histone octamers by salt dialysis, agarose gel electrophoresis, restriction enzyme protection, analytical ultracentrifugation, and electron microscopy to verify the correct number and positioning of the nucleosome cores are as described in (Bass et al., 2019). Additional reconstitutions with linker histone were performed by mixing reconstituted 183×12 core arrays with 1 mg/ml recombinant linker histone variant H1^0^ (New England Biolabs, cat.# M2501S) at a molar ratio of 0.7 molecule histone H1 per nucleosome in solution containing 500 mM NaCl, 10 mM HEPES, 0.1 mM EDTA and dialyzing against HNE buffer as described (Grigoryev et al., 2016).

### Mg^2+^-dependent chromatin folding and self-association assays

Native and reconstituted nucleosome arrays were dissolved in HNE buffer at final concentrations of 100 and 200 μg/mL for chromatin self-association assays and imaging analyses, respectively, and mixed with increasing concentrations of MgCl_2_ (Sigma Aldrich, cat. # M1028-1mL) and incubated for 20 minutes on ice. For some experiments, samples were incubated for up to 48 hrs. and used for further imaging or electrophoretic experiments.

The extent of chromatin self-association was analyzed using selective precipitation in magnesium as described in (Bass et al., 2019). The chromatin samples were incubated for 20 min. at different concentrations of MgCl_2_, and centrifuged at 12,000 rpm, 4°C, for 10 min in an Eppendorf 5810 R centrifuge with an F-34-6-38 rotor. Supernatants were collected and mixed with and equal volume of 12% glycerol, 40 mM EDTA, 2% SDS; the pellets were resuspended in 8% glycerol, 10 mM EDTA, 1% SDS. DNA from supernatants and pellets were analyzed on 1% agarose (Lonza cat. # 50000) gels in TAE buffer at 3 V/cm for 40 min. and stained with Ethidium Bromide or GelRedTM (VWR 89139-138). The percentage of DNA in the supernatant and pellet was determined by DNA band quantification using Image J software. The percentages of DNA at increasing MgCl2 concentration were input into Prism 9 for MacOS in order to interpolate a standard curve (Sigmoidal, 4PL, X is concentration) for both supernatant and pellet fractions. The IC50 values were used to determine the average concentration of MgCl_2_ at which 50% of the native or reconstituted chromatin was precipitated.

### PAD4-treatment of chromatin samples

To induce histone citrullination, the native or reconstituted chromatin samples (140 μg DNA/ml) in HNE were treated with 25 μg/ml human recombinant PAD4 enzyme (Sigma Aldrich cat # SRP0329 or Cayman Chemical cat. # 10500), 2.3 mM DTT, 1.9 mM CaCl_2_, 0.5 mM PMSF, and 25 mM NaHCO_3_. Control samples received the same treatment minus PAD4 or CaCl_2_. The PAD4-treated and control samples were incubated at 37°C for 1-2 hr. followed by 2 mM EGTA and placed on ice to stop the reaction. Histone citrullination was monitored by SDS-PAGE electrophoresis showing the downward mobility shifts of histone H3 and H4 and H1 due to its citrullination (Wang et al., 2004) and by Western blotting with antibodies against citrullinated histone H3 (Abcam ab5103).

### Electrophoretic and Western blotting techniques

For SDS-PAGE electrophoresis of control and PAD4 treated histones and band-shift analysis of citrullinated histones chromatin samples were dissolved in SDS-containing loading buffer and the electrophoresis was carried out in 18% acrylamide gels as described (Laemmli, 1970). The gels were stained with Brilliant blue R250 (FisherBiotech).

For SDS-PAGE electrophoresis followed by Western blotting, chromatin samples were dissolved in SDS-containing loading buffer and the electrophoresis was carried out in 8-16% mini-protean TGX Bio-Rad gels (Cat. # 4561105, Bio-Rad, Hercules, CA) gradient acrylamide gels as described (Laemmli, 1970). Proteins were transferred to Immobilon-P PVDF membrane (Cat. #PVH00010 Millipore, Bedford, MA) as described (Popova et al., 2009) and detected with primary antibodies against H3 citrullinated at Arginine 2, 8, and 17 (dil. 1:5.000, cat.# ab5103, Abcam, UK), secondary goat anti-rabbit HRP-conjugated antibodies (dil. 1:40.000, cat.# ab205718; Abcam, UK) and ECL prime Western detection kit (Cat. # RPN2232 GE Healthcare, UK)

For Triton-Acetic Acid-Urea (TAU) electrophoresis (Shechter et al., 2007) histones were acid-extracted from chromatin samples followed by precipitation with Trichloroacetic acid (TCA) as described (Sidoli et al., 2016). Extracted histones were resuspended in ddH_2_O and stored at -20°C until use. Concentration of extracted histones was approximated by serial dilution and SDS-PAGE electrophoresis in 18% acrylamide as before, and compared relative to a protein control. TAU sample buffer (0.9 M acetic acid, 16% glycerol, 5 mM DTT, 5.8 M urea, methyl green) was prepared fresh prior to TAU gel electrophoresis. Aliquots of stored histones were prepared at 6-10 μg/sample and dried by speed vacuum and resuspended in 10 uL of TAU sample buffer. Short TAU separating gel was prepared as described (Shechter et al., 2007) and placed in gel chamber with TAU running buffer (0.9 M acetic acid). Sample wells were flushed with running buffer using a syringe prior to loading samples. 10 uL of histones resuspended in sample buffer were loaded evenly into all lanes and run at 200 V constant for 2 hr with the electrodes reversed. Lanes without sample received 10 μ**L** of sample buffer. The gels were stained with Brilliant blue R250 (FisherBiotech).

For high-resolution DNA electrophoresis resolving the nucleosome repeats genomic DNA was purified from chromatin samples using the Wizard SV Gel and PCR Clean-Up System (Promega, catalog no. A9281) according to the manufacturer’s instructions. Samples were adjusted to a final concentration of 0.054 µg/µl with H2O and one-sixth the final volume of gel loading dye, no SDS (New England Biolabs, catalog no. B7025S). For size reference 1 kb and 100 bp DNA ladders (New England Biolabs, catalog no. N3232 and N3231) were combined and loaded onto the gel. The DNA samples were loaded to 0.49 ug and run on a 1.1% agarose gel (Sigma-Aldrich Type I-A, low EEO catalog no. A0169) in 1x Tris/acetic acid/EDTA (TAE) buffer (Bio-Rad, catalog no. 1610743) at 2.8 V/cm for 6 h 45 min with constant buffer recirculation. Gels were post-stained in GelRed (Biotium, catalog no. 41003) and digitally imaged. Gel analysis was performed using Image J software (National Institutes of Health) by measuring the relative migration distance (Rf) of the DNA standards and sample DNA bands. Rf values of the DNA standards were used to generate a standard curve of Rf against the log(Molecular weight) of the standards using Excel (Microsoft Excel for Office 365). The resulting linear equation was then used to determine the DNA length (bp) of each sample band. After determination of DNA length, the nucleosome repeat length was calculated as DNA length (bp) divided by the number of nucleosomes represented in each band.For DNP electrophoresis resolving the band shifts resulting from chromatin folding and self-association, we fixed the nucleosome arrays at 100-200 μg/mL with 0.4% formaldehyde for 10 min. on ice following 20 min incubation in HNE buffer containing various concentrations of MgCl_2_. The fixed arrays were then mixed with 1/5 total volume of 50% glycerol/HE (20 mM HEPES, 0.1 mM EDTA), 12 mM EDTA, and subjected to deoxynucleoprotein (DNP) electrophoresis in 1.0% Type IV agarose gel and run at 3 V/cm in HE buffer for 110 min. The agarose gels were stained by GelRed or ethidium bromide.

### Transmission Electron microscopy

For transmission electron microscopy, the samples resuspended in HNE containing appropriate concentrations of MgCl_2_, were fixed with 0.1 % glutaraldehyde for 16 hr and then dialyzed against HNE buffer. The dialyzed fixed samples were diluted to 1 μg/ml final concentration with 50mM NaCl, attached to carbon-coated and glow-discharged EM grids (EMS CF400C-Cu, Electron Microscopy Sciences), and stained with 2.0% uranyl acetate for negative staining (Woodcock and Horowitz, 1998). Bright-field EM imaging was conducted at 200 kV using JEM-2100 electron microscope (JEOL USA, Peabody, MA) equipped with 4k x 4k Ultrascan CCD camera (Gatan Inc. Warrendale, PA). EM images were collected at 60 - 80K nominal magnification.

### Cryo-Electron microscopy and tomographic reconstruction

Chromatin samples incubated for 20 min. without or with appropriate concentrations of MgCl_2_ were mixed with a suspension of 10 nm fiduciary gold particles (Sigma Aldrich # 741957), which were coated in bovine serum albumin to prevent clustering. 3 ul chromatin samples with a concentration of about 0.2 mg/ml DNA were applied to Quantifoil R2/2 200 mesh copper grids (EMS Q250-CR2). Vitrification was conducted by plunging into liquid ethane using our FEI Vitrobot Mk IV Grid Plunging System at 100% humidity, 4°C, and setting the blotting strength at 5, and blotting time at 3.5 sec.

Imaging of the vitrified samples was conducted on Titan Krios G3i 300 kV electron microscope, equipped with a K3 direct electron detector (Gatan, CA) at the Penn State Hershey cryo-EM core. We used Tomography-5.7.1. software (Thermo Fisher) for controlling data acquisition and collecting tilt-series. Cryo-EM tilt series (± 60°) were collected at 5° intervals in dose symmetric mode, at either 2.2 angstroms/pixel (defocus -6 um) or 1.7 angstroms/pixel (defocus -5 um), with zero-loss peak energy filtration through a 20-eV slit. Images were collected in 1x counting mode. Each tilt series had a cumulative dose of 120 electrons/Å^2^. Tilt series were aligned using fiducials in the IMOD software suite (https://bio3d.colorado.edu/imod/), CTF corrected and SIRT reconstructed. The chromatin samples, vitrification conditions, raw tilt series, and resulting cryotomograms are listed in the Suppl. Table 1.

### Centroid/axis/plane (CAP) modeling of nucleosome chain folding

The reconstructed un-binned tomograms were visualized and segmented into smaller subtomograms by IMOD/3dmod. Each volume was inverted using “newstack” to generate subtomograms with positive intensity corresponding to high density and filtered using IMOD command: “nad_eed_3d -n 30 -f -k 50” to reduce noise and enhance chromatin edges. The filtered subtomograms were exported into UCSF CHIMERA (RBVI, Univ. San Francisco, CA) for interactive visualization and analysis of nucleosome structures.

In CHIMERA, the filtered volumes were fitted with nucleosome core X-ray crystal structure (pdb 2CV5) semi-automatically to correspondent electron densities in the volume using the ‘fitmap’ command. The fitting of each nucleosome followed the local maximal electron density and thus was independent from the observer’s bias. After fitting, the nucleosomes were overlaid with centroids selecting the centers of masses, center-to-center axes, and nucleosome planes crossing the nucleosome at the dyad axis using the structure analysis ‘Axels/Planes/Centroids’ tool. For stereological analysis, each nucleosome in an array was numbered, if possible, starting from one end of the array and numbering consecutive nucleosomes to the other end of the array. Where linkers were not visible (such as in the tertiary chromatin structures), the nucleosomes were labeled based on their spatial proximity. The following measurements were recorded from such Centroid-Axial-Plane (CAP) models for each nucleosome (n) in an array: a) center-to-center distance D to the next nucleosome (n+1) in the array, b) center-to-center distance N to the nearest nucleosome (n_x_) in the 3D space, angle α between the two axes connecting each nucleosome with the previous one (n-1) and the next one (n+1) in a chain, angle β between the planes of consecutive nucleosomes n and n+1, and an angle *para* between the plane of each nucleosome (n) and the plane of the nearest nucleosome (n_x_) in the 3D space (see Fig 3 B). All distance and angle measurements for individual nucleosome arrays are included in the Suppl. Table 2.

The absolute nucleosome proximity values were obtained by subtracting the number of each nucleosome (n) from the number of its nearest nucleosome (n_x_) in the 3D space. The measured angle and plane values were analyzed statistically to determine the distribution profiles and average values that discriminate between the condensed and open nucleosomes arrays. Distance N and angle *para* were recorded for all nucleosomes. For some nucleosomes condensed by 0.75 mM Mg^2+^ (∼21% of all nucleosomes) and for all nucleosomes condensed at > 1 mM Mg^2+^, the linkers were not resolved and corresponding distances D and angles α and β were excluded from the statistical analysis. SDs were obtained from at least three tomograms and at least two independently cultured biological samples; *p-*values represent probability associated with a Student’s two-sample unequal variance t-test with a two-tailed distribution. The number of nucleosomes accounted for in each test is given in the corresponding figure legend.

