## Supplemental figures and legends for "Cryo-electron tomography reveals the multiplex anatomy of condensed native chromatin and its unfolding by histone citrullination" (from another version of this preprint: version 1, 2022-07-11)

**Supplementary Figure legends:**

**Suppl. Figure S1: Histone integrity and chromatin folding are not affected by an extended incubation in the presence of $Mg^{2+}$.**

(A) 18% SDS-PAGE gel stained by Coomassie R250 shows molecular mass markers (1) and histones of K562 fraction #10 chromatin incubated at the indicated concentration of $Mg^{2+}$ together with 10 nm fiduciary gold particles for 48 hrs. at +4°C (2 – 9).

(B, C) Reconstructed Cryo-ET tomograms of the K562 fraction #10 chromatin incubated at 0.75 mM $Mg^{2+}$ for 3 (B) and 16 (C) hrs. at +4oC before embedding in vitrified ice and imaging.

**Suppl. Figure S2: $Mg^{2+}$-dependent native chromatin folding is independent of the cell cycle.**

(A-C) FACS cell sorting profiles of K562 cells incubated at 37°C for 24 hours with normal supplemented media (A) and media supplemented with 0.5 mM mimosine (B) or 0.5 mM hydroxyurea (C). Cells were fixed with ethanol and stained with propidium iodide prior to analysis.

(D, E) $Mg^{2+}$-induced precipitation curves and [$Mg^{2+}$] at 50% sample precipitation determined for chromatin fractions #10 isolated from K562 cells incubated with normal supplemented media (control) and media supplemented with 0.5 mM mimosine or 0.5 mM hydroxyurea.

(F, G) Reconstructed Cryo-ET tomograms of the K562 fraction #10 isolated from K562 cells incubated with 0.5 mM HU (F) and 0.5 mM mimosine (G) and embedded in vitrified ice at the indicated concentration of $Mg^{2+}$ together with 10 nm fiduciary gold particles.

**Suppl. Figure S3: Nucleosome repeat length and Cryo-ET imaging of chromatin isolated from human HeLa and mouse retina cells.**

(A) Agarose gel shows DNA size markers (lane 1, 3, 5), DNA of native MNase-digested chromatin isolated from K562 cells (2), HeLa cells (4) and PN21 mouse retina cells (6).

(B) Linear equation of the standard curve from the migration distance (Rf) and log(MW) of DNA standards.

(C) Nucleosome repeat lengths (±S.D.) determined for K562, HeLa cells, and PN21 mouse retina cells.

(D) $Mg^{2+}$-induced 50% precipitation points determined for chromatin isolated from K562, HeLa cells, and PN21 mouse retina cells.

(E, F) Reconstructed Cryo-ET tomograms of the K562 fraction #10 isolated from chromatin isolated from HeLa cells €, and PN21 mouse retina cells (F) and embedded in vitrified ice at the indicated concentration of $Mg^{2+}$ together with 10 nm fiduciary gold particles. Scale bars: 50 nm.

**Suppl. Figure S4: Representative snapshots of CAP models of the unfolded and $Mg^{2+}$-condensed native nucleosome arrays isolated from K562 cells.**

(A) CAP models of the unfolded native nucleosome arrays vitrified at 0 mM $Mg^{2+}$.

(B) CAP models of the $Mg^{2+}$-condensed native nucleosome arrays vitrified at 0.75 mM $Mg^{2+}$. The models are grouped by the number of stacked nucleosomes per particle indicated at the top of each panel. The model numbers at the bottom of each panel correspond to those in Suppl. Table 2.

**Suppl. Figure S5: Comparison of internucleosomal distance and plane measurements for two independent batches of K562 cells vitrified in HNE buffer.**

(A) Box and whiskers plots of distances D and N measured for K562 batch 1 (blue) and K562 batch 2 (green) nucleosome arrays.

(B, C) Frequency distribution profiles of internucleosomal distances D (panel B) and N (panel C) measured for K562 batch 1 (blue) and K562 batch 2 (green) nucleosome arrays.

(D) Box and whiskers plots of angles $\alpha$, $\beta$, and *para* measured for K562 batch 1 (blue) and K562 batch 2 (green) nucleosome arrays.

(E–G) Two-dimensional plots showing distributions of the internucleosomal angles $\alpha$ (E), $\beta$ (F), and *para* (G) vs. the distance N for K562 batch 1 (blue crosses) and K562 batch 2 (green circles) nucleosome arrays.

(H) Average pairwise nucleosome proximities (per nucleosome array) determined for K562 batch 1 (blue columns) and K562 batch 2 (green columns).

**Suppl. Figure S6: Stereological data describing the path of nucleosome chain folding in the unfolded and $Mg^{2+}$-condensed native chromatin.**

(A) Frequency distribution profiles of distances D for arrays of nucleosomes vitrified at 0 mM $Mg^{2+}$ (green curves, n = 256) and arrays of nucleosomes vitrified at 0.75 mM $Mg^{2+}$ (blue curves, n = 402).

(B–D) Box and whisker plots of angles $\alpha$ (B), $\beta$ (C) and *para* (D) measured for total (100%), low 20% (low 20%) and top 20% arrays of nucleosomes vitrified at 0 mM $Mg^{2+}$ (green boxes, for 100% n = 220 ($\alpha$), 256 ($\beta$), 270 (*para*)) and arrays of nucleosomes vitrified at 0.75 mM $Mg^{2+}$ (blue boxes, for 100% n = 326 ($\alpha$), 396 ($\beta$), 507 (*para*)).

(E) Distances between the nucleosome number 1 (n1) and the subsequent nucleosomes (nX) were determined for the arrays of nucleosomes vitrified at 0 mM $Mg^{2+}$ (green circles) and arrays of nucleosomes vitrified at 0.75 mM $Mg^{2+}$ (blue squares).

(F) Average pairwise nucleosome proximities (per nucleosome array) determined for K562 nucleosome arrays vitrified at 0 mM $Mg^{2+}$ (green box, n = 32) and arrays of nucleosomes vitrified at 0.75 mM $Mg^{2+}$ (blue box, n = 48).

(G) Distribution of pairwise nucleosome interactions captured by *in situ* EMANIC in human HeLa cells. Data from (Grigoryev et al., 2016).

(H, I) Distributions of the ratio of nucleosomes with distance N < 8 nm per total nucleosomes (H) and average distance N in each array vitrified at 0 mM $Mg^{2+}$ (green line, n = 32) and arrays of nucleosomes vitrified at 0.75 mM $Mg^{2+}$ (blue line, n = 48).

**Suppl. Figure S7: Cryo-ET sections (Y-axis and Z-axis) show tight nucleosome packing inside the nucleosome condensates.**

Representative sections from Cryo- tomograms of K562 chromatin embedded in vitrified ice at 1.0 mM $Mg^{2+}$ (1-4, TS_5_2), 1.25 mM $Mg^{2+}$ (5-8, TS_6_2); and 2.0 mM $Mg^{2+}$ (9-12, TS_8_2); Sections 1, 5, and 9 were collected for the tomograms rotated perpendicularly to the Y-axis. The other sections were collected for the tomograms rotated perpendicularly to the Z-axis at three positions (I, II, and III) corresponding to those indicated by the dashed lines of the same color on the Y-axis section. Mag: 53,000-x; scale bars: 30 nm.

**Suppl. FigureS8: Cryo-ET imaging of the control and PAD4-treated native chromatin.**

(A-F): Cryo-tomograms showing control-treated K562 chromatin (A, C, E) and PAD4-treated K562 chromatin (B, D, F) vitrified in HNE buffer at 0 mM $Mg^{2+}$ (A, B), 0.75 mM $Mg^{2+}$ (C, D), and 1.0 mM $Mg^{2+}$ (E, F). Mag: 42,000-x; scale bars: 50 nm.

**Suppl. Figure S9: In the absence of $Mg^{2+}$, the unfolded native chromatin is not significantly altered by PAD4-dependent histone citrullination.**

(A) Box and whiskers plots showing average distances D and N measured for control-treated K562 chromatin (green boxes, n = 193 (D), 199 (N)) and PAD4-treated K562 chromatin (blue boxes, n = 189 (D), 194 (N)) at 0 mM $Mg^{2+}$.

(B) Box and whiskers plots showing average angles $\alpha$, $\beta$, and para measured for control-treated K562 chromatin (green boxes, n = 170 ($\alpha$), 193 ($\beta$), 197 (*para*)) and PAD4-treated K562 chromatin (blue boxes, n = 167 ($\alpha$), 189 ($\beta$), 194 (*para*)) at 0 mM $Mg^{2+}$.

(C) Average pairwise nucleosome proximities (per nucleosome array) determined for arrays of control-treated K562 chromatin (green bars, n = 22 arrays) and PAD4-treated K562 chromatin (blue bars, n = 22 arrays) at 0 mM $Mg^{2+}$.

(D-G) Frequency distribution profiles of distances N (panel H), angles $\alpha$ (I) and para (J) measured for arrays of control (green curves) and PAD4-treated (blue curves) nucleosome arrays vitrified at 0.75 mM $Mg^{2+}$.

(H–J) Two-dimensional plots showing distributions of the internucleosomal angles $\alpha$ (H), $\beta$ (I), and para (J) vs. the distance N for the control (green circles) and PAD4-treated (blue crosses) nucleosome arrays vitrified at 0.75 mM $Mg^{2+}$.

**Suppl. Figure S10: Stereological analysis of chromatin unfolding induced by histone citrullination in the $Mg^{2+}$-condensed native nucleosome arrays.**

(A-E) Box and whisker plots for distances D (A), N (B), and angles $\alpha$ (C), $\beta$ (D) and *para* (E) measured for total (100%), low 20% (low 20%) and top 20% arrays of nucleosomes from control (green boxes, for 100% n = 209 (D), 302 (N), 162 ($\alpha$), 210 ($\beta$), 302 (para)) and PAD4-treated K562 chromatin (blue boxes, for 100% n = 162 (D), 215 (N), 130 ($\alpha$), 161 ($\beta$), 215 (para)) vitrified at 0.75 mM $Mg^{2+}$.

(F-G) Frequency distribution profiles of distances D (F) and angles $\beta$ (G) measured for arrays of control (green curves n = 209 (D), 210 ($\beta$)) and PAD4-treated (blue curves n = 162 (D), 161 ($\beta$)) K562 chromatin vitrified at 0.75 mM $Mg^{2+}$.

(H) Distribution of pairwise nucleosome proximities measured for control (green bars n = 178) and PAD4-treated (blue bars n = 142) K562 chromatin vitrified at 0.75 mM $Mg^{2+}$.

**Suppl. Figure S11: Stereological analysis of control and PAD4-treated 183x12+H1 reconstituted nucleosome arrays at 0.75 mM $Mg^{2+}$.**

(A) Box and whisker plots of internucleosomal distances D and N calculated for control (green) and PAD4-treated (blue) reconstituted +183x12+H1 nucleosome arrays vitrified at 0.75 mM $Mg^{2+}$. Control samples: n = 204 (D), n = 230 (N); PAD4-treated samples: n = 150 (D), n = 158 (N).

(B) Box and whisker plots for angles $\alpha$, $\beta$ and *para* measured for control-treated (green bars n = 197 ($\alpha$), 217 ($\beta$), 230 (para)) and PAD4-treated K562 chromatin (blue bars n = 135 ($\alpha$), 150 ($\beta$), 158 (para)) 183x12+H1 reconstituted nucleosome arrays at 0.75 mM $Mg^{2+}$.

(C-D) Frequency distribution profiles of angles $\beta$ (C) and *para* (D) measured for arrays of control (green curves n = 217 ($\beta$), 230 (para)) and PAD4-treated (blue curves n = 150 ($\beta$), 158 (para)) 183x12+H1 reconstituted nucleosome arrays at 0.75 mM $Mg^{2+}$.

(E) Average pairwise nucleosome proximities (per nucleosome array) determined for control-treated (green box, n = 20 arrays) and PAD4-treated (blue box n = 14 arrays) 183x12+H1 reconstituted nucleosome arrays vitrified at 0.75 mM $Mg^{2+}$.

(F) Distances between the nucleosome number 1 (n1) and the subsequent nucleosomes (nX) were determined for control-treated (green circles n = 15) and PAD4-treated (blue circles n = 11) 183x12+H1 reconstituted nucleosome arrays vitrified at 0.75 mM $Mg^{2+}$.

Supplementary Figure S1

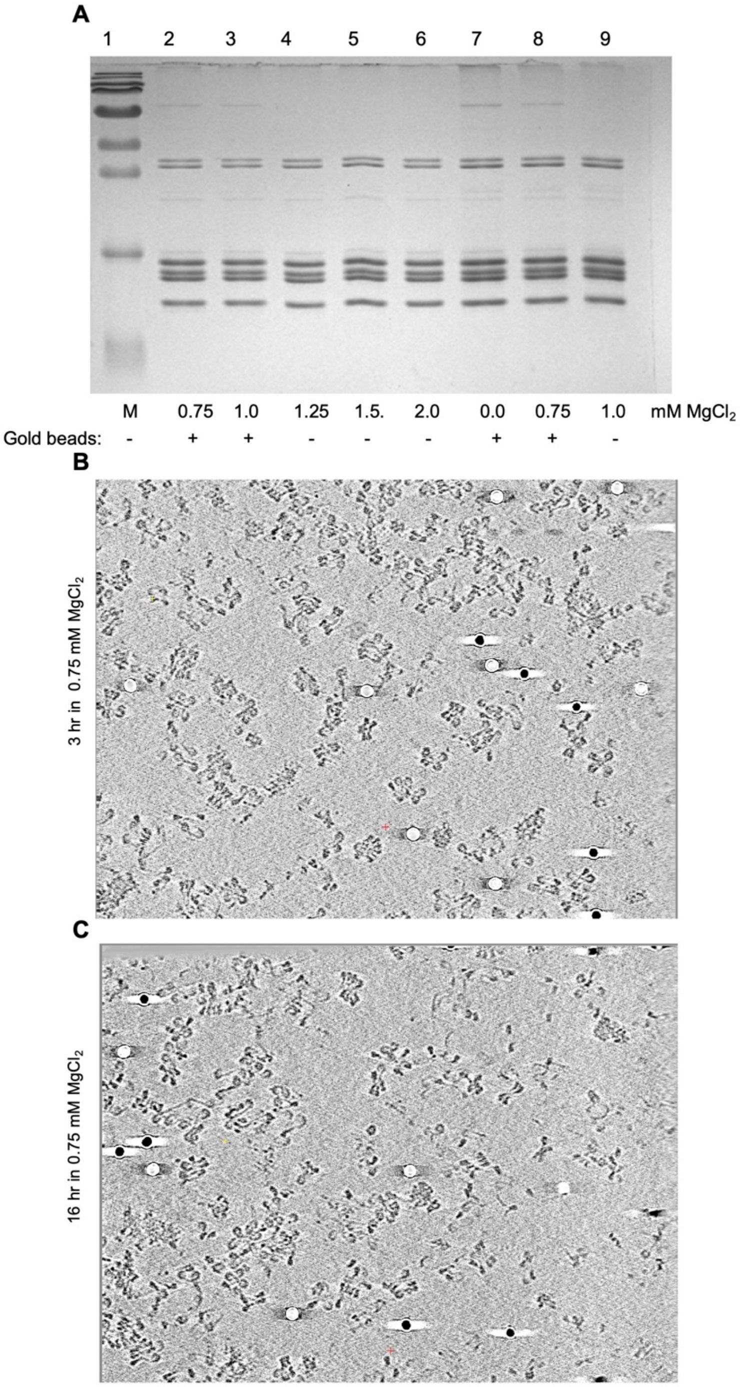

Supplementary Figure S2

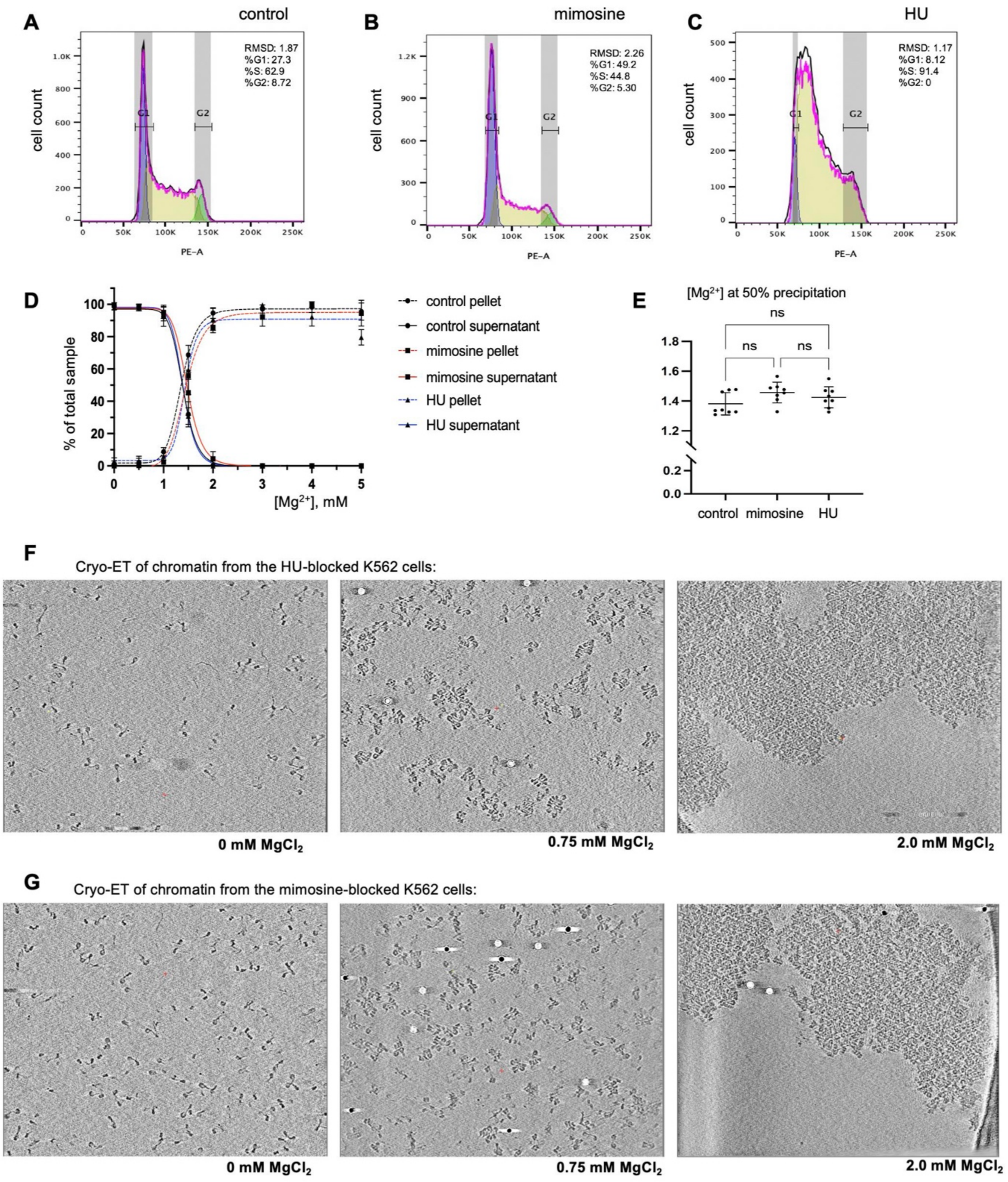

Supplementary Figure S3

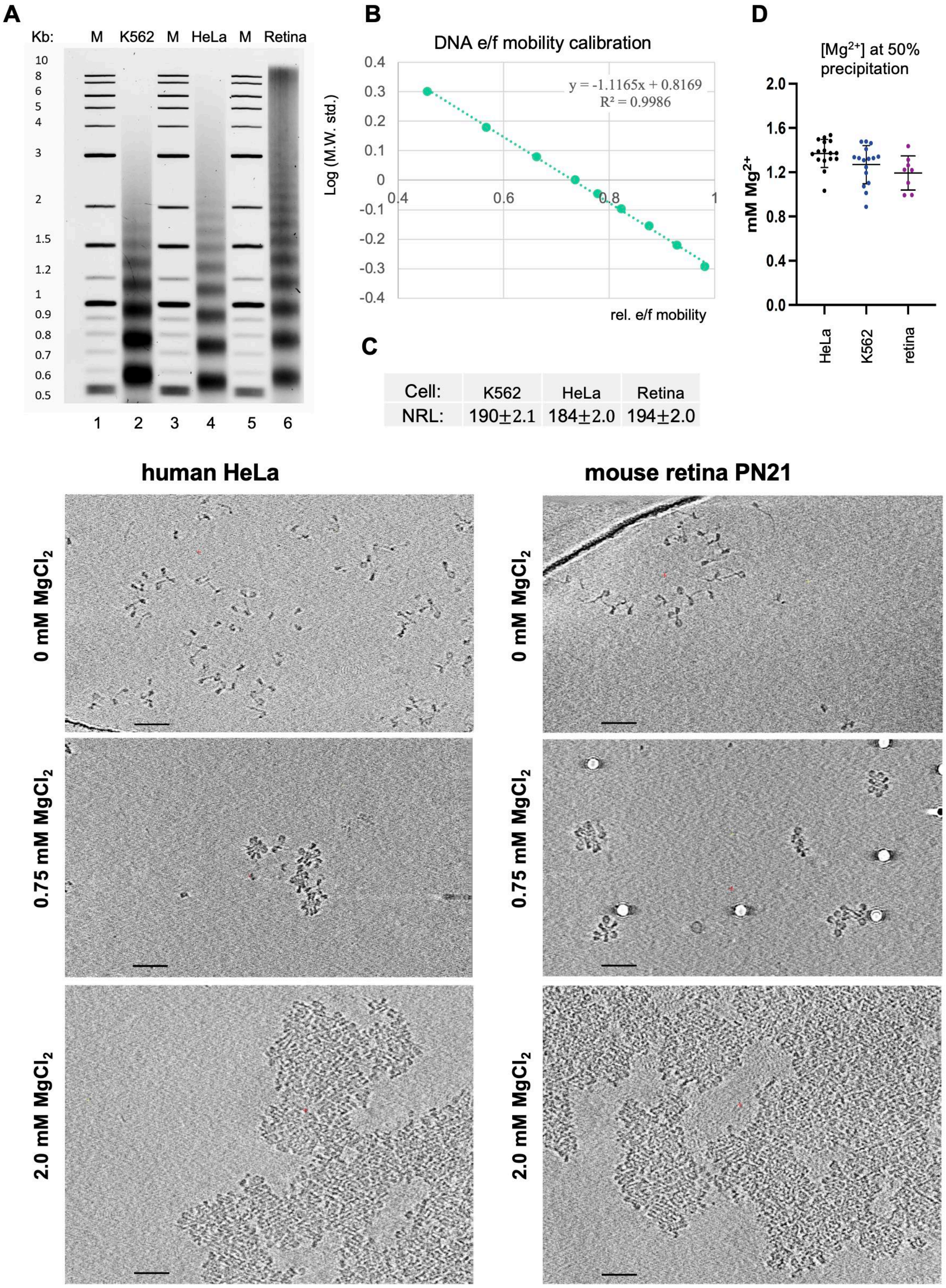

Supplementary Figure S4 A

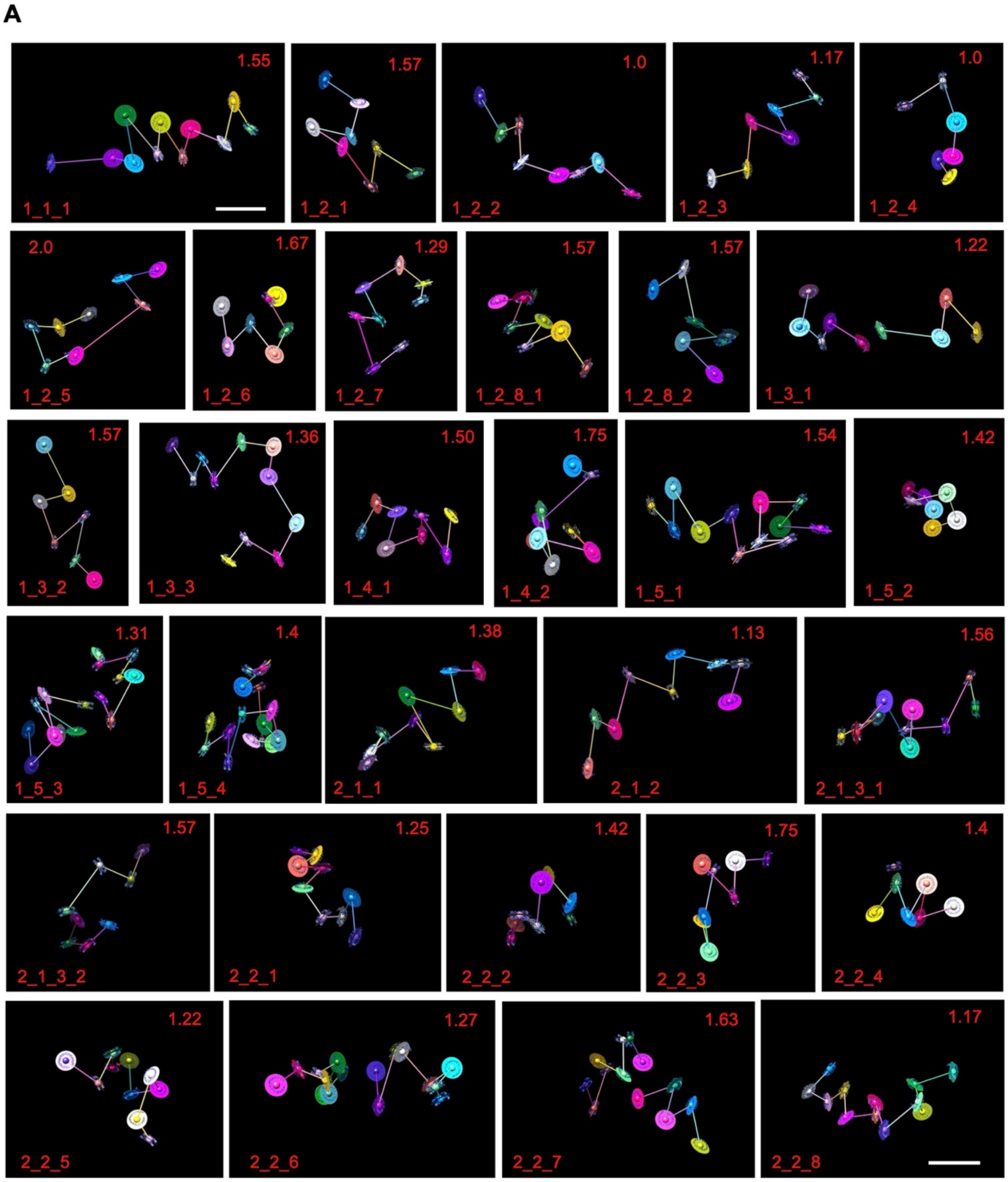

Supplementary Figure S4 B

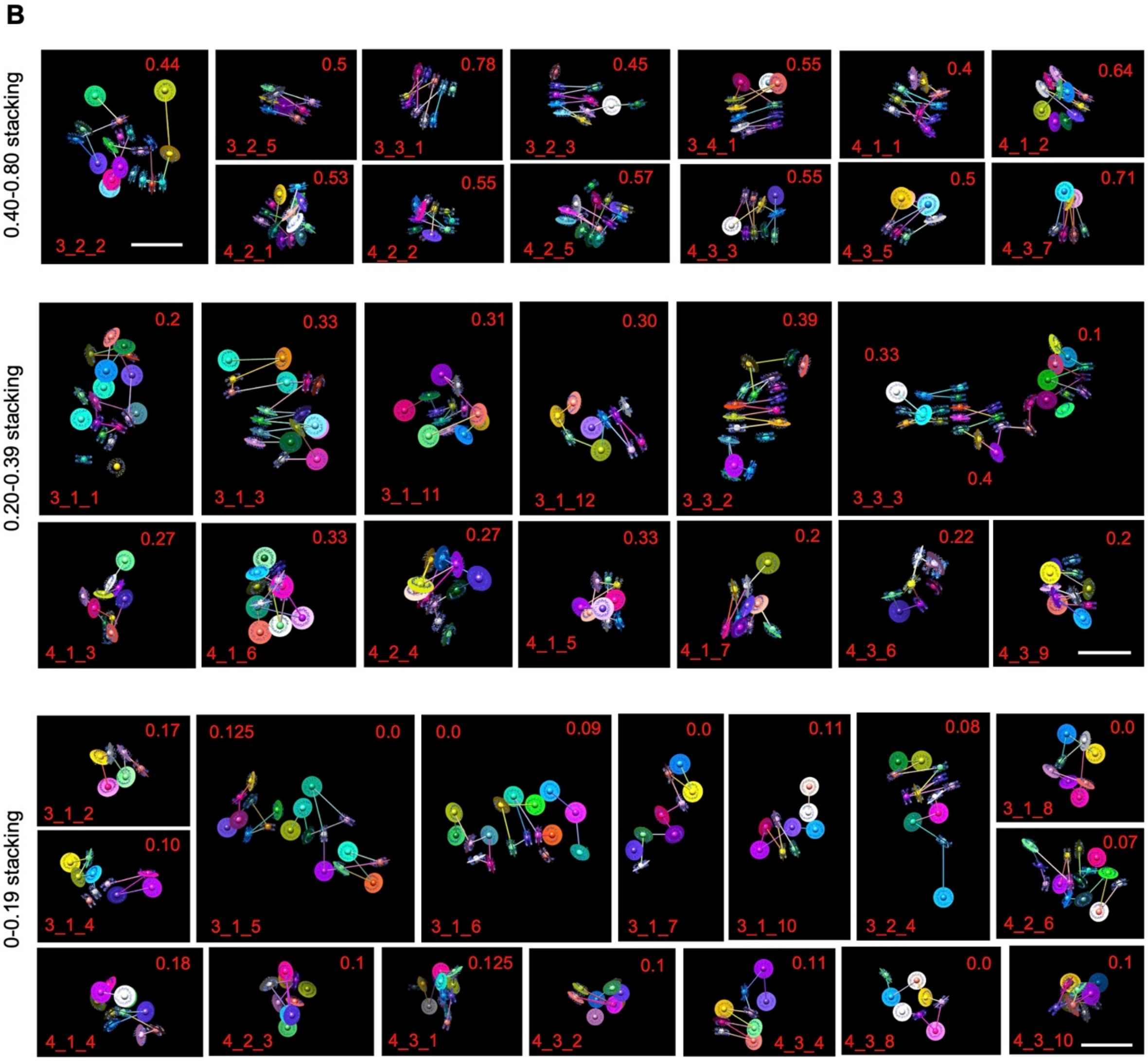

Supplementary Figure S5

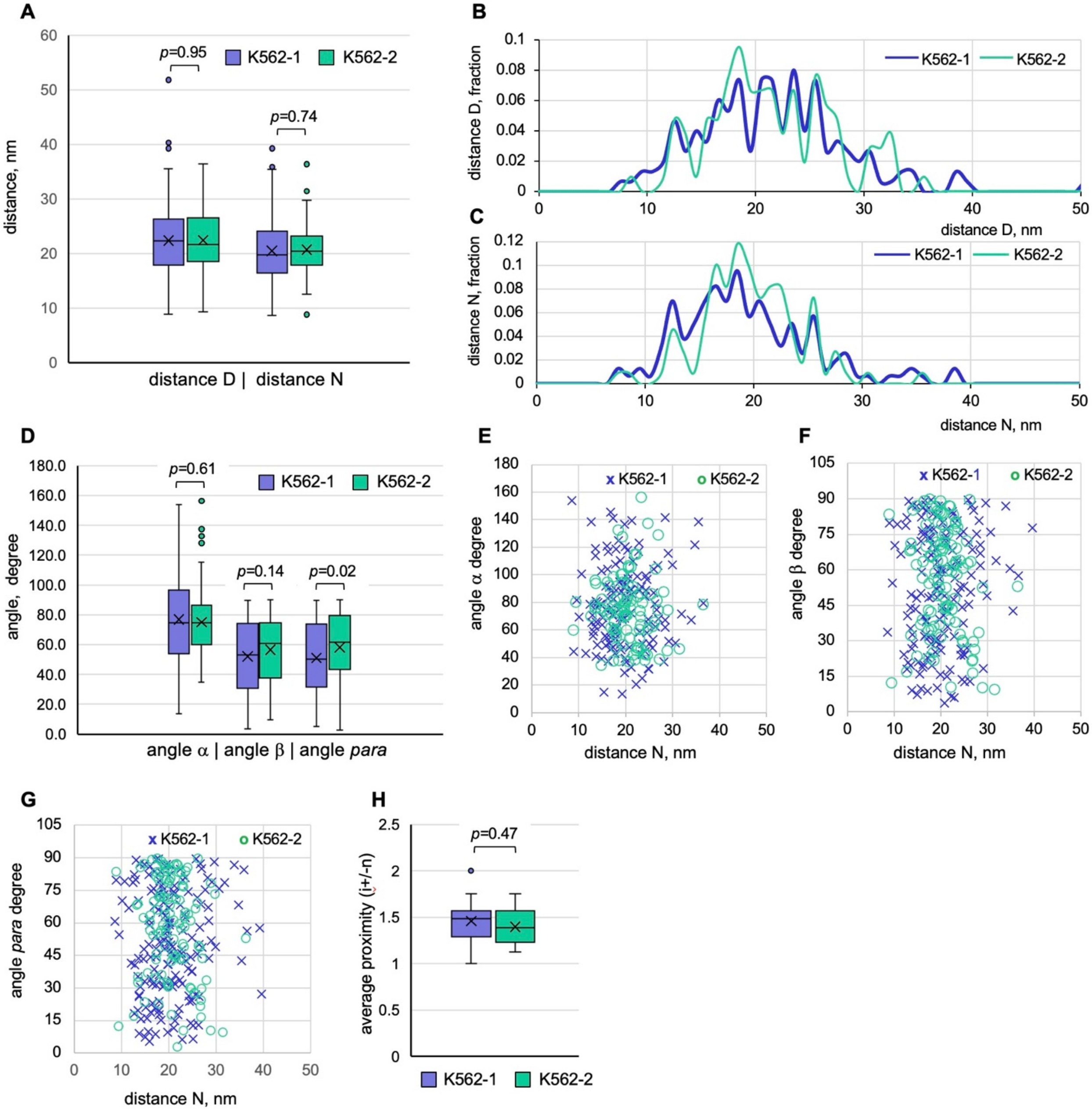

Supplementary Figure S6

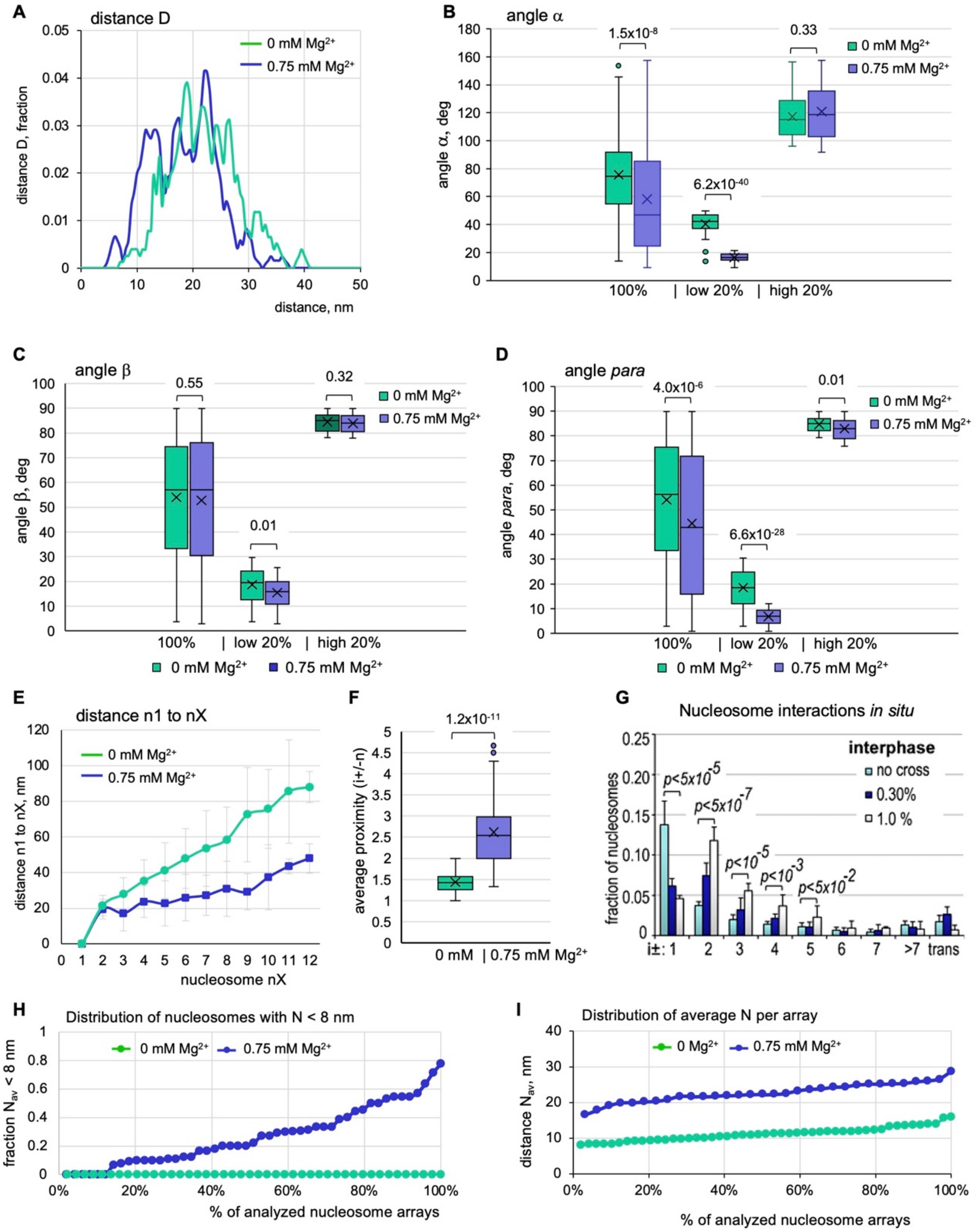

Supplementary Figure S7

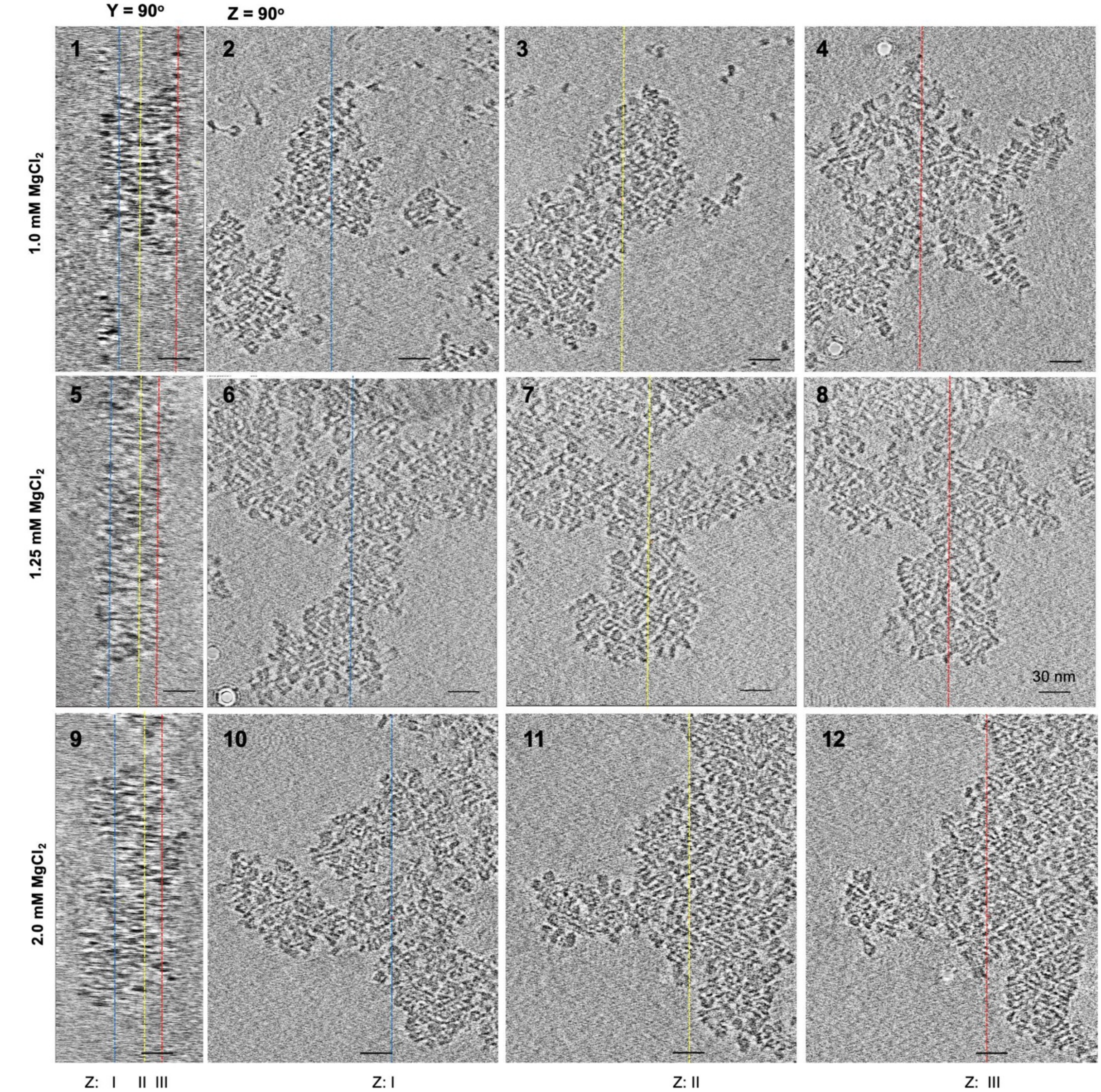

Supplementary Figure S8

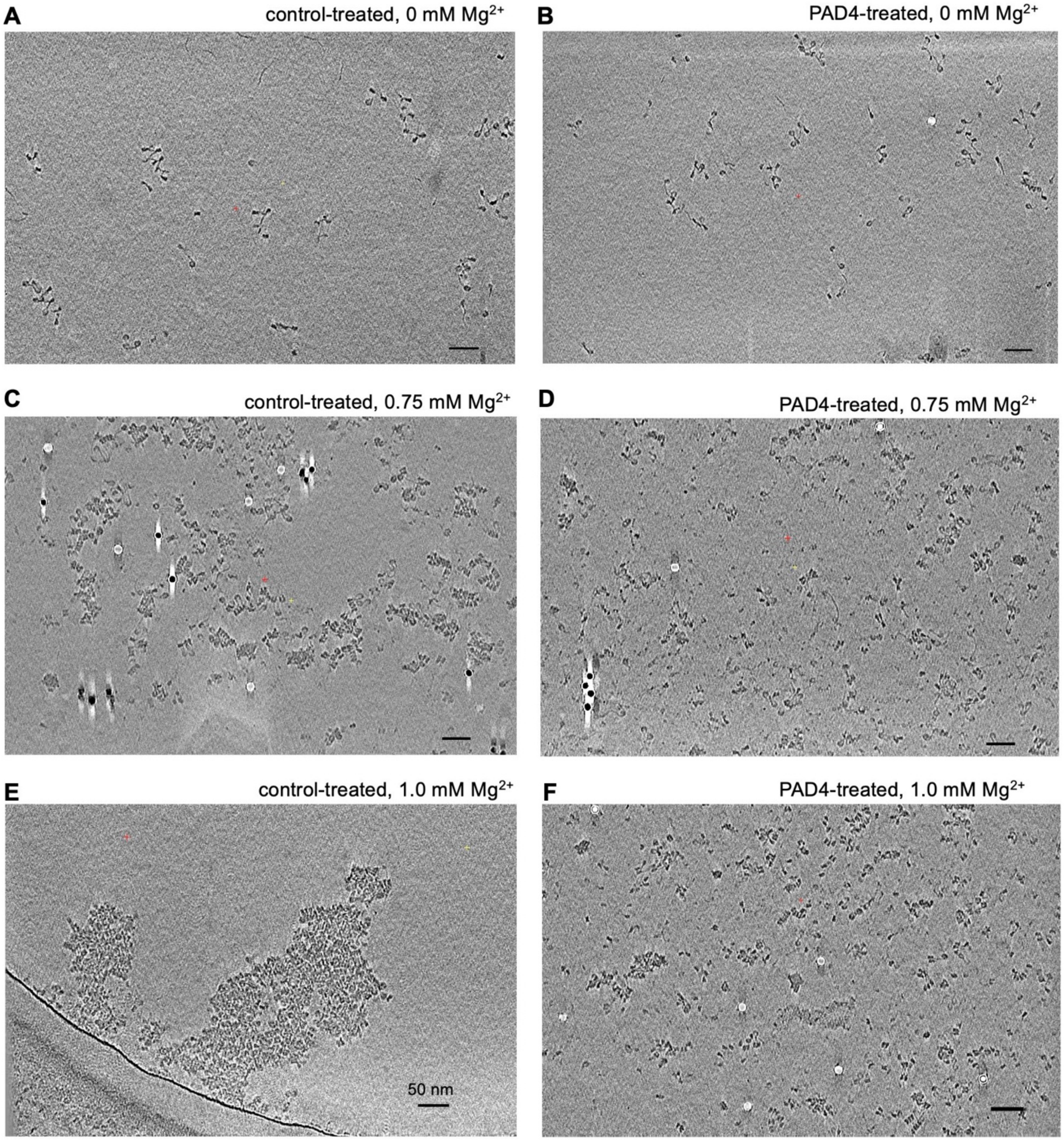

Supplementary Figure S9

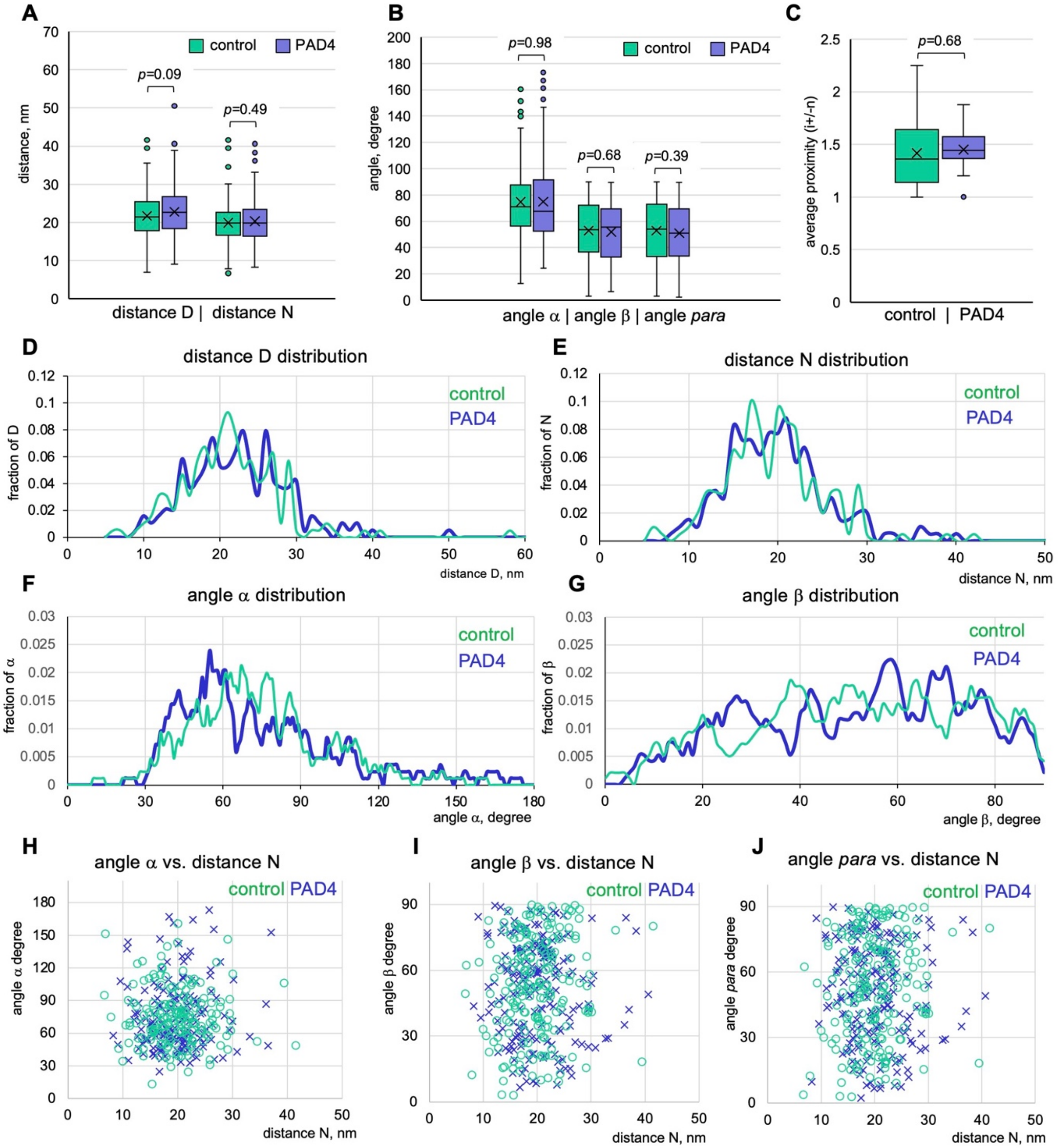

Supplementary Figure S10

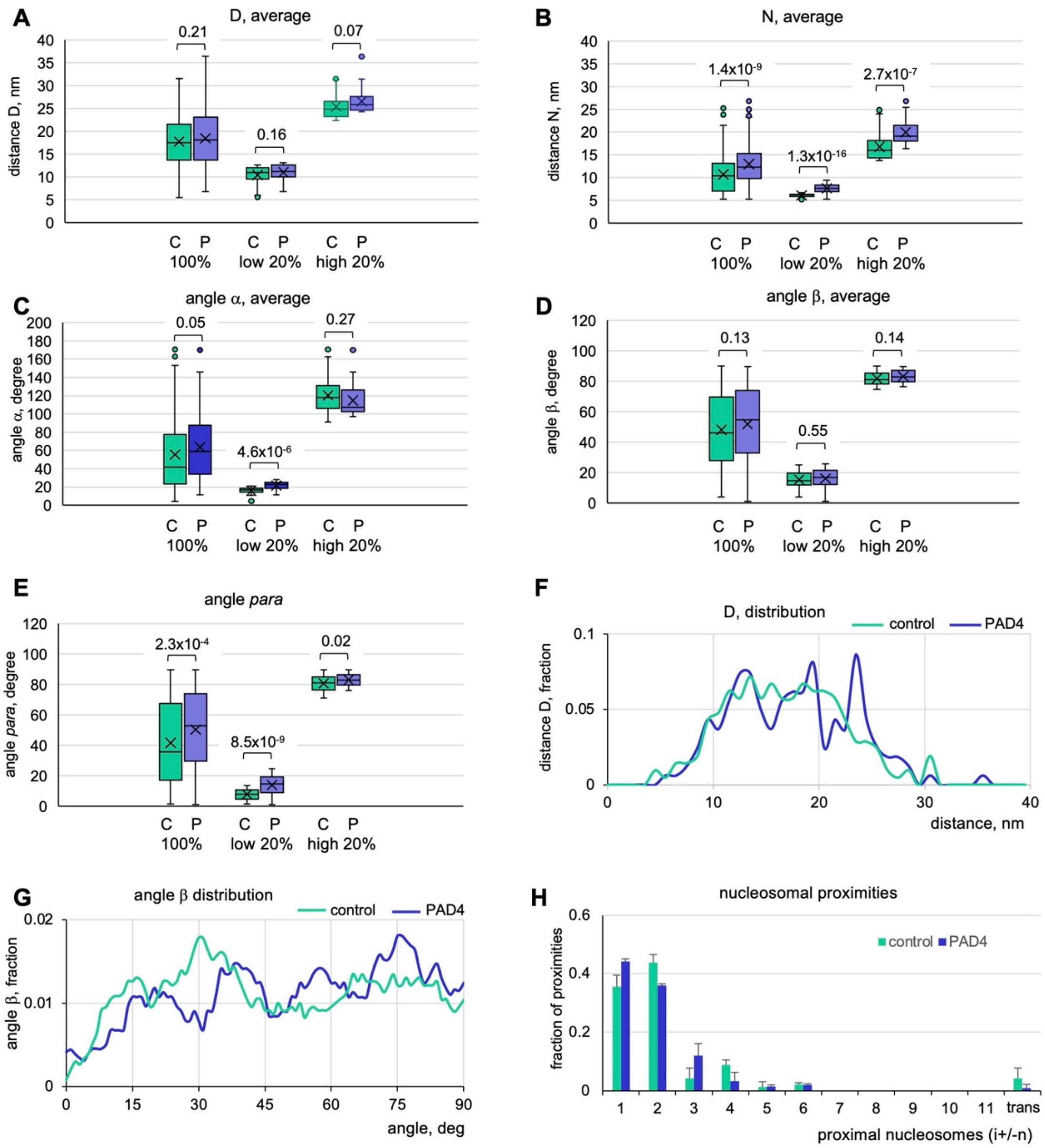

Supplementary Figure S11

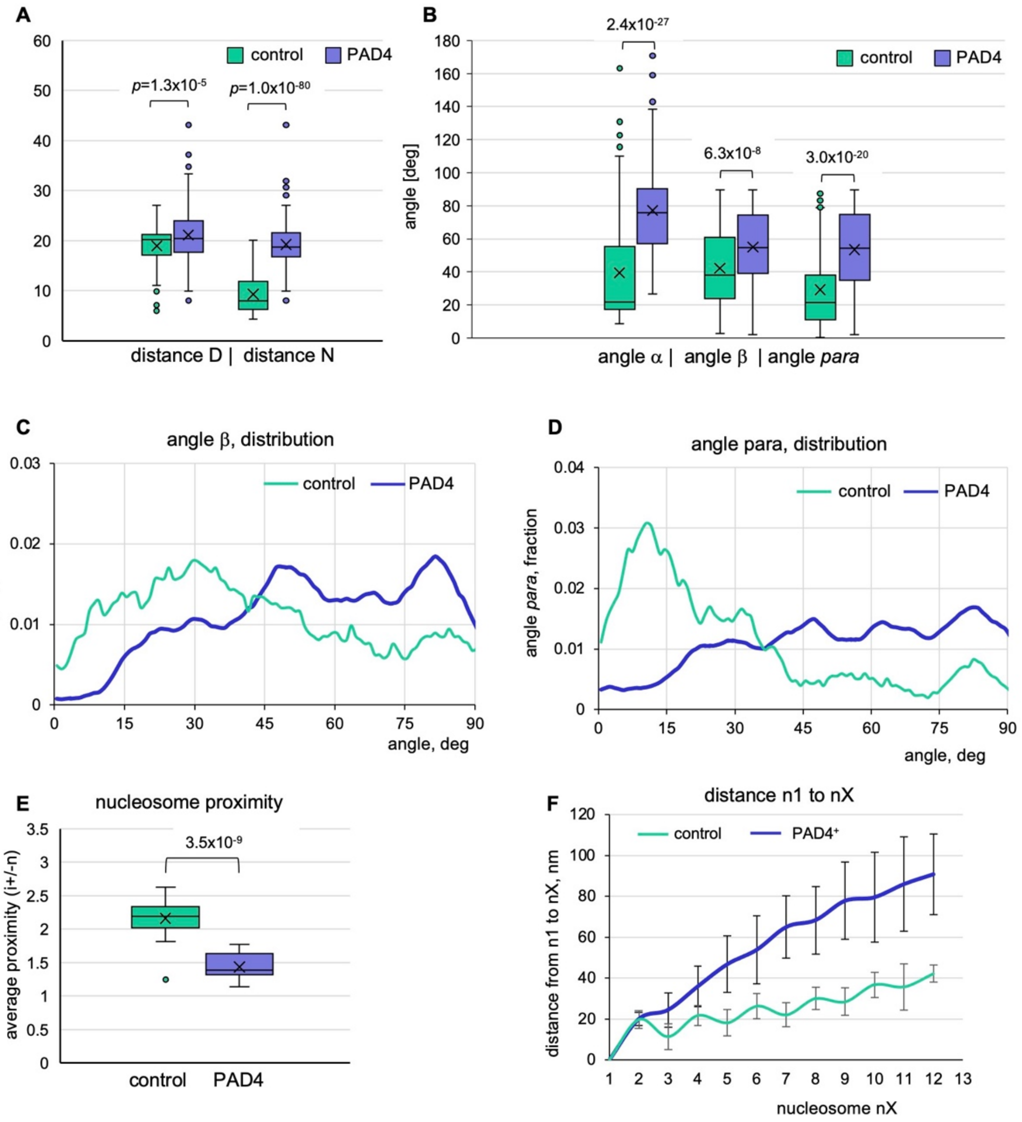
